# Increased complexity and incomplete conservation of human distal lung regeneration

**DOI:** 10.64898/2026.07.27.741036

**Authors:** Hsiao-Yun Liu, Camilla Predella, Juan A. Torres, Elisabeth Schönweiler, Tania A. Thimraj, Aaron D. Springer, Songjingyi Liang, John W. Murray, Anjali Saqi, Jeremy Worley, Ya-Wen Chen, N. Valerio Dorrello, Hans-Willem Snoeck

## Abstract

As curative treatments for many lung diseases remain elusive, deeper insight into human lung regeneration is essential. In mice, surfactant-producing type 2 alveolar epithelial (AT2) cells regenerate type 1 (AT1) cells, which mediate gas exchange, and are replenished from airway secretory subsets after severe injury. Here we show computationally that in humans and ferrets, but not in mice or swine, AT2-independent trajectories from terminal airway subsets and from proximal airway basal cells to AT1 cells operate, which were validated in human organoid and human-rat xenograft models, respectively, and are disrupted in a disease-specific fashion. These trajectories include a TGFβ and Hippo-regulated physiological equivalent of profibrotic aberrant basaloid cells present in diseased lungs that are precursors for an injury-associated AT1 subset. The incomplete conservation of lung regeneration has important implications for the pathogenesis of lung disease and for determining the most appropriate models to study normal and aberrant human lung repair.

## INTRODUCTION

Animal models have allowed groundbreaking discoveries that contributed to our understanding of human development and biology. However, while the initial stages of vertebrate development are highly conserved, later developmental stages are more divergent to accommodate different organismal sizes, life spans, ecological niches and positions in the food web. Possibly as a result of this divergence, faithful recapitulation of several human diseases in animal models has been challenging. This is also true in the lung, where the development of disease-modifying treatments for intractable diseases has been hindered by the lack of reliable mouse models.^1^ A prime example is idiopathic pulmonary fibrosis (IPF), an intractable familial or sporadic fibrotic interstitial lung disease (ILD)^2^ that has not been faithfully reproduced in mice. The classical model, administration of bleomycin (BLM), causes a transient, diffuse, inflammatory fibrosis,^3^ rather than the non-inflammatory, focal and progressive fibrosis, called ‘usual interstitial pneumonia’ (UIP), that is pathognomonic for IPF.^4^ Although mutations in genes involved in telomere maintenance are the most frequent cause of familial IPF,^5^ late generation telomerase-deficient mice with critically shortened telomeres do not show fibrosis or increased BLM susceptibility.^6^ Furthermore, mice with conditional deletion of the telomere maintenance gene, shelterin (*Trf2*), succumb to acute lung injury and not to progressive fibrosis after BLM exposure.^7^ Proteotoxic surfactant protein mutations are rare causes of familial IPF.^8^ Transgenic expression of mutant human surfactant proteins in mice can lead to perinatal mortality,^9–11^ acute lung injury when expressed at high levels,^12^ spontaneous fibrosis, though not with the UIP pattern,^12,13^ or increased BLM-induced fibrosis,^14^ overall a spectrum of phenotypes more similar to childhood ILD. Mice with mutations in Hermansky-Pudlak Syndrome (HPS) genes that cause and early-onset, IPF-like disease with nearly 100% penetrance,^15,16^ show increased BLM sensitivity but not spontaneous fibrosis.^17–20^ Finally, the bronchitis component of chronic obstructive pulmonary disease (COPD) is not recapitulated in cigarette smoke-exposed mice.^21^

The difficulty to recapitulate human lung diseases in mice may be due at least in part to incomplete conservation of mechanisms involved in lung regeneration. While most organs have dedicated stem and progenitor compartments, regeneration in the distal lung relies on facultative stem cells endowed with extensive plasticity. In mice, surfactant-producing type 2 alveolar epithelial (AT2) cells can self-renew and convert to AT1 cells, the flat alveolar cells that mediate gas exchange.^22–25^ This transition occurs predominantly through a KRT8^+^ ‘alveolar differentiation intermediate’ (ADI) population.^26–28^ ADI cells express senescence and extracellular matrix (ECM) markers ^29^. AT2 cells can also regenerate from cells at the bronchioalveolar junction^30,31^ and, after severe injury, from subsets of airway secretory cells (SCs).^32–37^ After severe injury, TRP63^+^KRT5^+^ cells resembling airway basal cells (BCs) arise from SC-like progenitors that can organize as ‘pods’ (KRT5-pods).^38–41^ Their significance is unclear as they have been associated with dysplastic, maladaptive repair^39,42–47^ or with alveolar regeneration.^36,40^ Similar structures were also observed in human acute lung injury and were interpreted as peribronchiolar squamous metaplasia.^48,49^

Differences in distal lung anatomy between humans and mice may impart distinct regenerative and pathogenic mechanisms, however. In contrast to rodents, BCs also reside in intrapulmonary airways in humans.^50^ In mice, distal airways connect directly to alveolar sacs. Lungs in many larger mammals contain a zone of terminal and respiratory bronchioles (TRBs), consisting of small airways and alveolar structures that connects airways with alveolar acini.^51^ While present in cats, dogs, primates and ferrets, TRBs are much less developed in swine, sheep, horses and cattle, however ^52^. TRBs in humans are lined by cell types (collectively called ‘TRBCs’ here) subsets of which share expression of SCGB3A2 with SCs and of SFTPB with AT2 cells. A role in regeneration is suggested by the accumulation of TRBC subsets in injured lungs.^53–55^

In several lung diseases, including IPF,^48,54^ COPD^56^ and fibrotic ILD secondary to the autoimmune disease, systemic sclerosis (SSc-ILD),^57^ *KRT17^+^KRT5^-^*‘aberrant basaloid cells’ (aBCs) were identified. aBCs may be profibrotic by virtue of their expression of ECM components and location near fibroblastic foci in IPF^54,58,59^ and are deemed orthologous to ADI cells,^29^ although the latter do not express KRT17. Rare cells of similar phenotype have been identified in normal lungs,^54^ however, suggesting a physiological role. Studies using precision-cut lung slices^60^ and human pluripotent stem cells (hPSC)-derived AT2 (iAT2) cells^61^ suggested that human aBCs, similar to mouse ADI cells, can arise from AT2 cells. However, our previous studies in organoids mutant for the *HPS1* gene^62^ indicated that aBCs can also be derived directly from TRBCs, implicating intrinsic TRBC dysfunction in IPF pathogenesis.^1,62^

Gaining insight into regenerative trajectories in the human lung and into the role of dysfunction of human lung progenitor populations in the pathogenesis of lung disease requires the use of human cell-based models and humanized animal models to complement animal models. Here we show using comparative trajectory analysis, organoid approaches and a novel human-rat xenograft model that in the human lung AT1 cells also regenerate from TRBCs and from BCs through trajectories that include aBC-like cells. These trajectories are absent in mice and pigs but conserved in ferrets, and are disrupted in a disease-specific fashion. The extraordinary plasticity of the human lung and the incomplete conservation of mechanisms involved in distal lung regeneration has profound implications for our understanding of human lung disease and for the development of more appropriate cell-based and animal models.

## RESULTS

### Comparative trajectory analysis of human, pig, ferret and mouse lungs

We re-analyzed publicly available scRNAseq datasets, subsetted on non-ciliated epithelial cells, from lungs in mice, pigs, ferrets and humans **(Supplementary Table 1)**. We focused on two mouse datasets using BLM^63,64^ and one using influenza^27^ as injury models. We also included a dataset from ferrets treated with two doses of BLM^65^ and one from pigs in homeostasis.^66^ As human datasets, we used the integrated IPF and COPD datasets from the Human Lung Cell Atlas (HLCA),^67^ a single nucleus RNAseq dataset from patients who succumbed to Covid,^68^ and a dataset from patients with SSc-ILD.^57^

In homeostasis, ferret and human distal lung contained, in addition to AT1 and AT2 cells, TRBCs, BCs and SCs. In contrast to human TRBCs, however, *SCGB1A1*, not *SCGB3A2*, was the predominant SC marker in ferret *SFTPB^+^* TRBCs **(Fig. 1, Fig. S1)**, suggesting reversal of expression patterns or misannotation in the ferret genome. Consistent with the absence of TRBs in these species,^52^ TRBCs were not identified in pig and mouse lungs, where only SCs (mouse) or SCs and BCs (pig) were present. In pigs a small *KRT8^+^ITGB6^+^* subset was present that clustered between AT1 and AT2 cells, which we labeled as ADI cells. In the human homeostatic dataset, we identified a small cluster expressing some *KRT17* and *ITGB6*, but not *KRT5* or *TP63*, a phenotype consistent with aBC-like cells **(Fig. 1, Fig. S1)**. After injury, subsets coalesced, suggesting regenerative plasticity. In mice, a *KRT8^+^* ADI population appeared, confirming previous reports **(Fig. 1, Fig. S1)**.^26–28^ In human and ferret injury datasets, airway subsets became more preponderant, with profound depletion of AT1 cells in IPF and SSc-ILD **(Fig. 1, Fig. S1)**. In IPF, AT1 cells became embedded among airway cells **(Fig. 1, Fig. S1)**. In all human injury datasets, an aBC-like (Covid) or aBC (IPF, SSc-ILD, COPD) population was prominently present. In BLM-treated ferrets, a population of *KRT8^+^ITGB6^+^* cells appeared, which we classified as ‘aBC-like’ **(Fig. 1, Fig. S1)**. We note that *KRT17* is not annotated in the ferret genome and could not be used to identify aBC-like cells.

**Figure 1.**
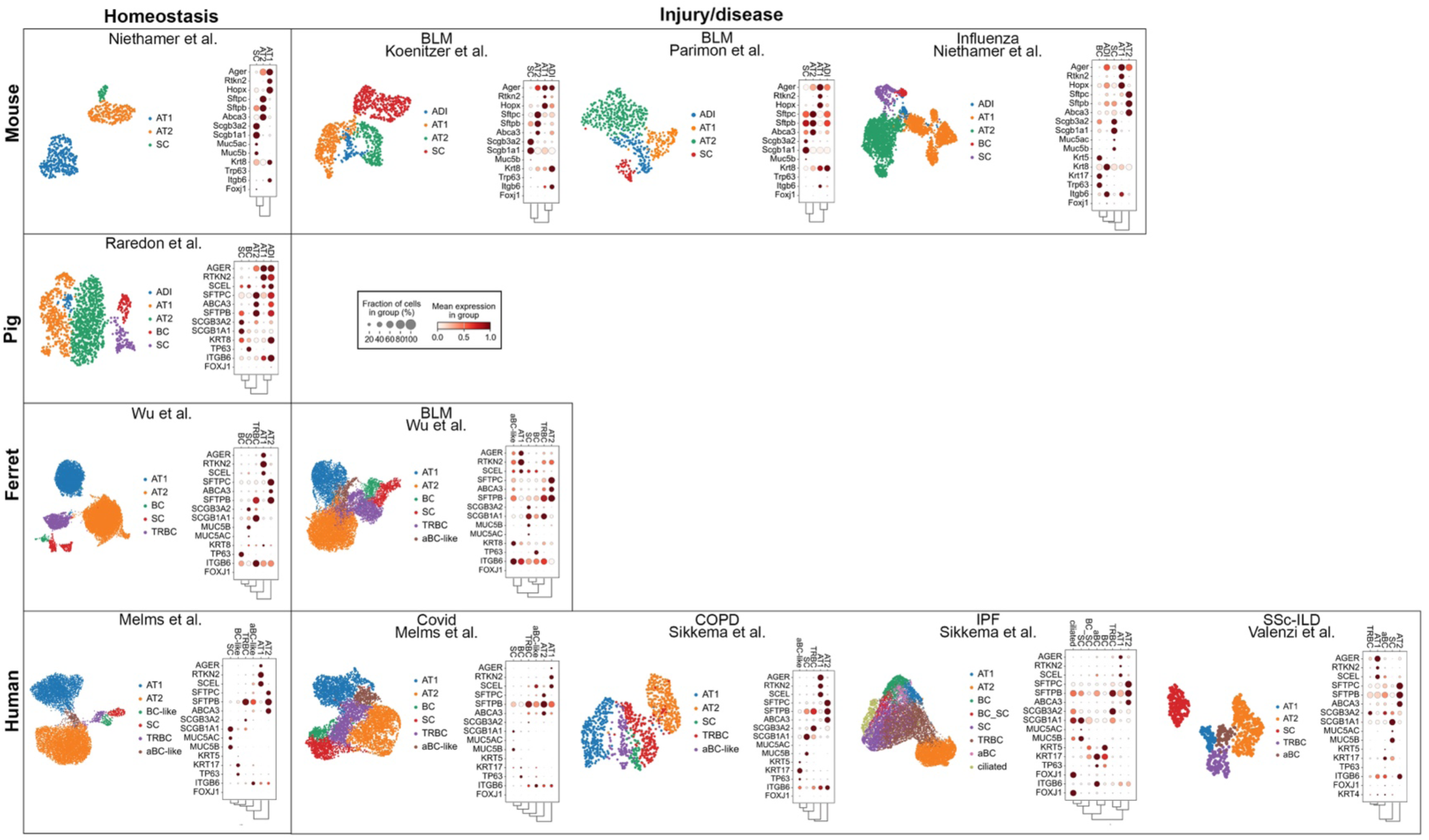
In silico comparison of mouse, human, pig and ferret lung regeneration. UMAP plots of cell type assignment based on canonical markers shown in dotplots in mice, ferret and human lung in homeostasis and after injury or disease.

We next performed trajectory analysis using partition-based graph abstraction (PAGA)^69^ **(Fig. 2A)** and modeled transitions between select subsets by diffusion pseudotime (DPT)^70^ **(Fig. 2B)**. PAGA indicated only sparse cell fate transitions in mice and ferrets in homeostasis, which were therefore not included in this analysis. In the mouse influenza and BLM models and in homeostatic pig lungs, PAGA placed ADI cells between AT1 and AT2 cells **(Fig. 2A)**, with a continuous increase in DPT along AT2>AT2 and AT2>ADI>AT1 trajectories **(Fig. 2B)**, consistent with published lineage tracing studies in the mouse.^22,24–26,28,29,36,71^ In the influenza model, where injury was presumably more severe, SCs and downstream *Krt5^+^Trp63^+^* BC-like cells provided input into the ADI subset as well. Trajectories in the pig and mouse lung are therefore highly conserved, although pig lungs appear less quiescent in homeostasis.

**Figure 2.**
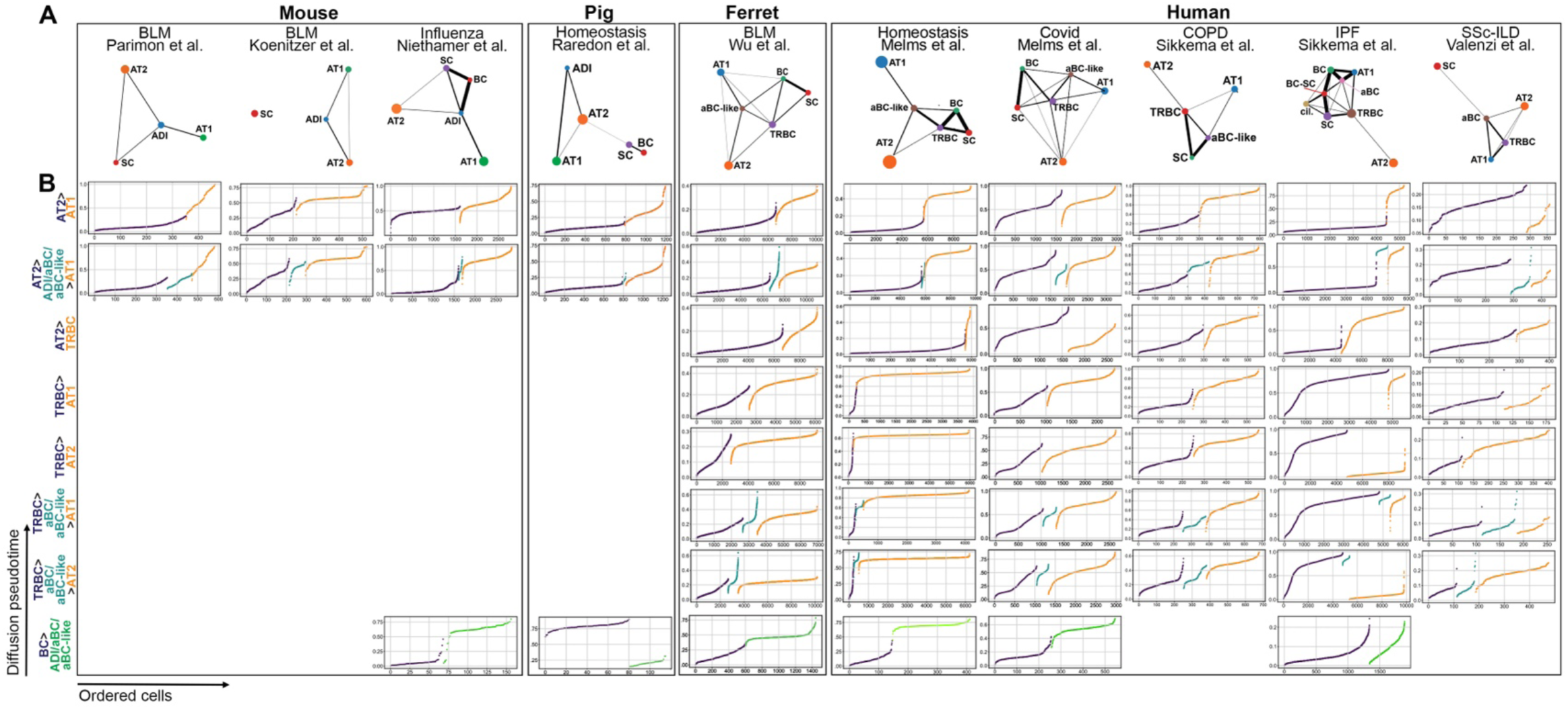
Trajectory analysis of mouse, human and ferret lung regeneration. **A.** PAGA trajectory analysis in datasets from Fig. 1. **B.** DPT analysis along trajectories indicated on the left.

In all human datasets and in the BLM-treated ferret dataset, PAGA revealed dense connectivity between aBCs and aBC-like, TRBCs, BCs, AT1 and AT2 cells **(Fig. 2A)** with TRBCs as a central node, suggesting an important role in regeneration. In human homeostatic lungs, smooth DPT transitions along AT2>AT1, TRBC>AT1 and AT2>TRBC and TRBC>AT2 trajectories **(Fig. 2B)** indicate extensive plasticity. Inclusion of aBC-like cells in these trajectories showed discontinuity in aBC-like>AT2 but not in aBC-like>AT1 trajectories, indicating that aBC-like cells are dedicated AT1 progenitors. In BLM-treated ferrets, similar trajectories were identified, although inclusion of aBC-like cells always showed the highest DPT in aBC-like cells, suggesting a differentiation block in this cell population. In the human injury datasets aBCs and aBC-like cells showed closer connectivity to airway cells and AT1 cells than to AT2 cells, which consistently appeared as a more isolated population. This finding may indicate a more minor role of AT2 cells in regeneration. Specific trajectories were disrupted in disease states. In Covid, trajectories emanating from AT2 cells were discontinuous, suggesting either a more important role of TRBC-driven regeneration or AT2 dysfunction. In COPD and IPF, the AT2>AT1 transition appeared preserved, while DPT curves along trajectories emanating from TRBCs were disrupted, especially in IPF where the highest DPT was always reached in TRBCs and aBCs, suggesting TRBC dysfunction in both diseases, especially in IPF. In SSc-ILD, on the other hand, TRBC>AT2 and AT2>TRBC transitions were preserved, while all trajectories towards AT1 cells were disrupted. These data indicate a specific AT1 differentiation defect, irrespective of the precursor cells, in SSc-ILD. Finally, in all datasets, PAGA suggests existence of two trajectories from TRBCs to AT1 cells, a direct trajectory and a trajectory through aBC-like cells.

Given the consistent connectivity of ADI cells, aBC-like cells and aBCs on one hand and BCs on the other, we next modeled these trajectories from BCs by DPT analysis **(Fig. 2B, bottom panels)**. BC>ADI transition appeared unlikely in mice and pigs. Human homeostatic and Covid lungs and ferret BLM-treated lungs, however, showed smooth DPT transitions from BCs to aBC-like cells. BCs may therefore provide a major input into the aBC-like population, and thus into the AT1 subset.

Taken together, *in silico* analysis suggests conservation of the AT2>AT1 transition with additional regenerative trajectories in humans and ferrets emanating from TRBCs and BCs, where BCs are entirely AT1-biased. aBC-like are consistently present as intermediates in all trajectories towards AT1 cells. The trajectories inferred in mice are consistent with fate-mapping studies,^24–26,28,29,35–37,39^ thus serving as experimentally validated positive controls. We next tested the veracity of the inferred trajectories in humans using organoid modeling for TRBCs and xenotransplantation for BCs.

### A KRT17^+^ injury-associated AT1 differentiation pathway

We previously reported the generation of TRBC-like cells from hPSCs (embryonic (ESC) and induced pluripotent (iPSC) stem cells) and their differentiation into AT1 cells.^62^ Expandable spheres, called ‘induced respiratory airway progenitors’ (iRAPs), are generated from lung organoids (LOs)^72,73^ and match TRBC populations as defined by Murthy et al. **(Fig. 3A)**.^55^ Consistent with computational analysis **(Fig. 2)**, iRAPs possess AT1 potential. Differentiation into induced AT1 (iAT1) cells **(Fig. 3A)** entails first a transitional stage, in which WNT signaling is withdrawn and TGFβ signaling inhibited (TGFβi) and where most cells express *SCGB3A2* together with AT1 or AT2 markers, followed by plating in 2D in the presence of TGFβi and BMP4 resulting in cultures where ∼95% of the cells express only AT1 markers.

**Figure 3.**
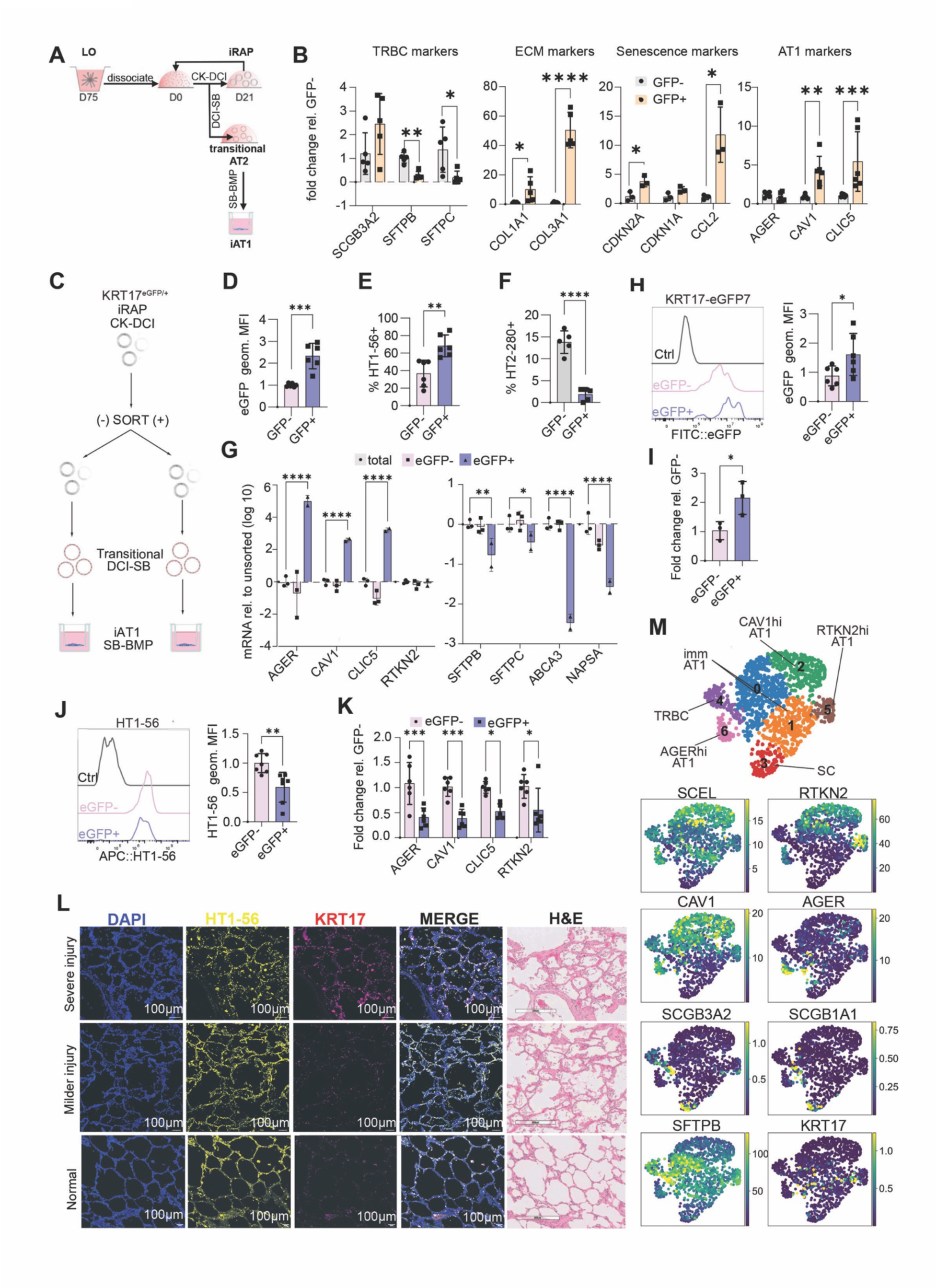
KRT17^+^ AT1 lineage from iRAPs. **A.** Schematic representation of iRAP and iAT1 differentiation protocol. **B.** Expression of TRBC markers (n=5), collagens (n=5), senescence (n=3), and AT1 (n=6) markers in KRT17-eGFP iRAPs normalized to eGFP-. *p<0.05, **p< 0.01, ****p<0.0001. unpaired two-sided Student’s t-test. **C.** Schematic of generation of transitional and iAT1 cells. **D-F.** eGFP geometric MFI normalized to eGFP^-^**(D),** percent HT1-56^+^ **(E)** and HT2-280^+^ **(F)** cells among transitional AT2 cells generated from KRT17-eGFP^+^ and KRT17-eGFP^-^iRAPs. **p < 0.01, ***p < 0.001, ****p < 0.0001, unpaired two-sided Student’s t-test, n=6. **G.** mRNA expression of AT1 (left) and AT2 markers (right) in sorted and unsorted transitional AT2 cells. *p<0.05, **p<0.01 and ****p<0.0001, two-way ANOVA with Tukey’s multiple comparison test, n=3 for total and eGFP^-^, n=2 for eGFP^+^. **H.** Representative flow cytometry histogram (left) and MFI (right) of eGFP expression normalized to eGFP^-^in iAT1 cells derived from KRT17-eGFP^-^and KRT17-eGFP^+^ iRAPs. *p<0.05, unpaired two-sided Student’s t-test, n=6. **I.** *KRT17* mRNA in iAT1 cells derived from KRT17-eGFP^-^and KRT17-eGFP^+^ iRAPs. *p < 0.05, unpaired two-sided Student’s t-test, n=3. **J.** Representative flow cytometry histogram (left) and quantification of HT1-56 MFI (right) normalized to eGFP^-^in iAT1 cells derived from KRT17-eGFP^-^and KRT17-eGFP^+^ iRAPs. **p<0.01, unpaired two-sided Student’s t-test, n=6. **K.** mRNA expression of AT1 markers in iAT1 cells derived from KRT17-eGFP^-^and KRT17-eGFP^+^ iRAPs. *p<0.05, ***p<0.001, two-way ANOVA with Šidák multiple comparison test, n=6. **L.** IF for KRT17 in human lungs with diffuse alveolar damage. **M.** UMAP of clusters among AT1 cells in Covid lungs and feature plots for AT1, AT2 and TRBC markers and *KRT17*.

To examine the generation of aBC-like cells from iRAPs, we generated a KRT17-eGFP reporter ESC line **(Fig. S2A)**. iRAPs derived from this line contained a KRT17-eGFP^+^ subset **(Fig. S2B,C)** that was quiescent **(Fig. S2D)**, and showed reduced expression of the TRBC markers, *SFTPB and SFTPC,* as well as increased expression of the aBC markers*, COL1A1, COL3A1, CDKN2A, CDKN2B and CCL2,* and of the AT1 markers, *CAV1 and CLIC5* **(Fig. 3B)**. Bulk RNAseq comparing KRT17-eGFP^+^ and KRT17-eGFP^-^iRAPs yielded 71 significantly differentially expressed genes that included aBC markers (*CCL2* and *CDKN2B*) and AT1 markers (*SCEL* and *CLIC5*) **(Fig. S2E)**. Pathway analysis showed that intermediate filament organization, keratinization and skin development were significantly induced in the KRT17-eGFP^+^ fraction **(Fig. S2F)**. The expression of both aBC and AT1 markers in the KRT17-eGFP^+^ fraction is consistent with computational inference **(Fig. 2)** suggesting that aBC-like are AT1 progenitors.

We therefore next determined the AT1 potential of purified KRT17-eGFP^+^ and KRT17-eGFP^-^ iRAPs.^62^ At the first, transitional stage **(Fig. 3C)**, cells generated from KRT17-eGFP^+^ iRAPs expressed more KRT17-eGFP than those derived from KRT17-eGFP^-^iRAPs (**Fig. 3D, Fig. S2G)**. Furthermore, cells generated from KRT17-eGFP^+^ iRAPs were enriched in HT1-56^+^ (a human AT1-specific surface marker)^74^ cells **(Fig. 3E, Fig. S2G)** and depleted of HT2-280^+^ (a human AT2-specific surface marker)^75^ cells **(Fig. 3F, Fig. S2G)**. They also expressed more mRNAs encoding AT1 markers and fewer mRNAs encoding the AT2 markers than those derived from KRT17-eGFP^-^iRAPs **(Fig. 3G)**. These observations suggest more rapid commitment to the AT1 fate. In the next stage, the 2D iAT1 stage **(Fig. 3C),** iAT1 cells generated from KRT17-eGFP^+^ iRAPs expressed more KRT17-eGFP **(Fig. 3H)** and *KRT17* mRNA **(Fig. 3I)** than iAT1 cells generated from KRT17-eGFP^-^iRAPs. However, the former expressed less HT1-56 **(Fig. 3J)** and lower levels of mRNAs encoding AT1 markers **(Fig. 3K)** than the latter, suggesting generation of a distinct AT1 or less mature subset. KRT17-eGFP^-^ iRAPs also generated iAT1 cells expressing KRT17, though at lower levels than KRT17-eGFP^+^ iRAPs **(Fig. 3H)**. KRT17-eGFP^-^ cells can therefore transition to KRT17^+^ iAT1 cells as well, albeit less efficiently. These *in vitro* studies suggest that aBC-like cells are intermediates in a KRT17^+^ AT1 differentiation pathway that commits to an AT1 fate more rapidly, but may generate a distinct or immature AT1 subset.

As KRT17^+^ AT1 cells had never been identified, we verified their presence in injured lungs. Immunofluorescent staining (IF) revealed locoregional presence of KRT17^+^ AT1 cells in severely injured areas of lungs from patients with acute respiratory distress syndrome, but not in normal controls **(Fig. 3L)**. To define which AT1 cells expressed KRT17, we subsetted AT1 cells in the Covid dataset (insufficient AT1 cells were identified in the other datasets subsets for meaningful analysis). This analysis revealed three clusters each characterized by high expression of either *CAV1*, *RTKN2* or *AGER*, suggesting AT1 heterogeneity after injury **(Fig. 3M)**. Small clusters of contaminating SCs (*SCGB3A2^+^SCGB1A1^+^SFTPB^+^*) and TRBCs (*SCGB3A2^+^SCGB1A1^+^SFTPB^-^)* were also detected, of which the significance is unclear. The remaining cells expressed *SCEL* and lower levels of the aforementioned AT1 markers **(Fig. 3M)**. KRT17 was expressed in a fraction of these AT1 cells and was also expressed in a subset of RTKN2^hi^ AT1 cells **(Fig. 3M)**.

Collectively, our observations indicate the existence of an injury-associated AT1 differentiation pathway from TRBCs through aBC-like cells that is at least transiently marked by KRT17, shows more rapid AT1 commitment *in vitro*, and yields AT1 cells with a distinct expression pattern of canonical AT1 markers. The existence of two AT1 differentiation trajectories from TRBCs is consistent with computational inference in human lung injury datasets **(Fig. 2)**.

### aBC-like cells are driven by Hippo inhibition and TGFβ

To gain mechanistic insight into the dual nature of KRT17-eGFP^+^ iRAPs, which share features of IPF-associated aBCs and of AT1 progenitors, we exposed iRAPs to TGFβ, which drives fibrosis,^76^ and to Hippo inhibition, which promotes AT1 regeneration.^77,78^

Hippo signaling phosphorylates the kinases, LATS1 and LATS2, which in turn phosphorylate the homologues YAP and TAZ, resulting in cytoplasmic retention. Inhibition of LATS induces nuclear translocation of YAP and TAZ where they associate with the transcriptional regulators, TEAD1-4, which play an essential role in AT1 regeneration after injury.^77,79,79–82^ IF revealed that ∼60% of nuclear (n)YAP^+^ iRAPs expressed KRT17-eGFP^+^, whereas only ∼10% of nYAP^-^iRAPs did **(Fig. 4A,B)**. nYAP is therefore largely associated with KRT17-eGFP^+^ iRAPs. In the presence of a small molecule LATS inhibitor (LATSi)^83^ virtually all cells became nYAP^+^, whereas TGFβ1 abrogated nYAP expression **(Fig. 4A,C)**. Both LATSi **(Fig. 4A,D)** and TGFβ1 **(Fig. 4A,E)** increased the fraction of KRT17-eGFP^+^ cells, however. Conversely, small molecule inhibition of the YAP-TEAD interaction (TEADi) **(Fig. 4D)** and TGFβi **(Fig. 4E)** each educed the KRT17-eGFP^+^ fraction.

**Figure 4.**
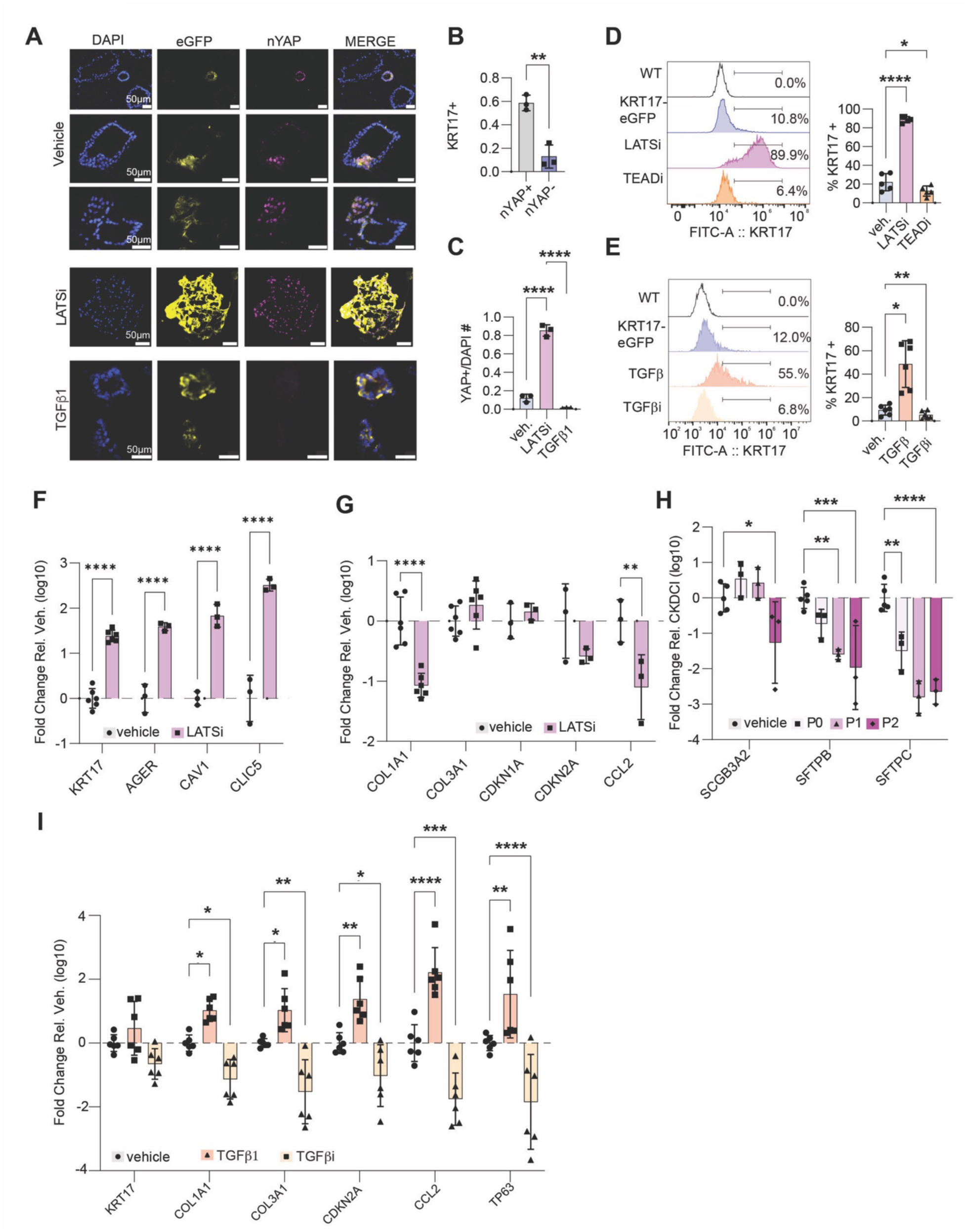
Regulation of KRT17 by TGFβ and Hippo signaling. **A.** IF representative of 3 biological replicates of KRT17-eGFP iRAPs in vehicle, LATSi and TGFβ1-treated iRAPs. **B.** Fraction KRT17^+^ cells in nYap^+^ and nYAP^-^iRAPs. **p < 0.01, unpaired two-sided Student’s t-test, n=3. **C.** Fraction of nYAP^+^ cells after addition of LATSi or TGFβ1. ****p < 0.0001, one-way ANOVA with Tukey’s multiple comparison test, n=3. **D.** Representative flow cytometry histogram (left) and quantification of fraction of *KRT17*^+^ cells (right) in the presence of LATSi or TEADi. *p < 0.05, ****p < 0.0001, one-way ANOVA with Holm-Šídák’s multiple comparisons test, n=5. **E.** Representative flow cytometry histograms (left) and quantification of fraction of *KRT17*^+^ cells (right) in the presence of TGFβ1 or TGFβi. *p < 0.05, **p < 0.01, one-way ANOVA with Dunnett’s multiple comparison test, n=6. **F.** mRNA expression of AT1 markers in LATSi and vehicle treated iRAPs. **p < 0.01, ***p < 0.001, ****p < 0.0001, one-way ANOVA with Tukey’s multiple comparison test, n=6 (*KRT17*) and n=3 (remaining). **G.** mRNA expression of aBC markers in LATSi and vehicle treated iRAPs. *p < 0.05, **p < 0.01, one-way ANOVA with Tukey’s multiple comparison test, n=6 (*COL1A1* and *COL3A1*) and n=3 (remaining). **H.** mRNA expression of TRBC markers in LATSi treated iRAPs over multiple passages. *p < 0.05, **p < 0.01, ***p < 0.001,****p < 0.0001, two-way ANOVA with Dunnett’s multiple comparison test, n=3. **I.** mRNA expression of aBC markers in TGFβ1 and TGFβi treated cells. *p < 0.05, **p < 0.01, ***p < 0.001, ****p < 0.0001, two-way ANOVA with Dunnett’s multiple comparison test, n=6.

While both LATSi and TGFβ signaling induced KRT17, the nature of the KRT17-eGFP^+^ cells differed. LATSi enhanced cellular expansion **(Fig. S3A)**, increased expression of AT1 markers **(Fig. 4F),** decreased expression of some aBC markers (*COL1A1*, *CDKN2A*, *CCL2)* **(Fig. 4G)**, and, over several passages, decreased expression of TRBC markers **(Fig. 4H)**. LATSi changed iRAP morphology from individual spheres to a ‘bunch of grapes’ **(Fig. S3B)**. Removing LATSi restored morphology to individual spheres **(Fig. S3B)**, reduced KRT17-eGFP and HT1-56 expression **(Fig. S3C,D)**, and restored AT1 and TRBC marker expression over several passages **(Fig. S3E)**. Hippo inhibition in iRAPs therefore reversibly induces and expands AT1-biased aBC-like cells from TRBCs.

In contrast to LATSi, TGFβ1 halted proliferation **(Fig. S3F)**. TGFβ1 increased, while TGFβi decreased, expression of canonical aBC markers **(Fig. 4I)**. TGFβ1 also reduced expression of TRBC markers **(Fig. S3G)**. Furthermore, *TGFB1* and *TGFBR1* mRNAs were elevated in KRT17-eGFP^+^ cells **(Fig. S3H),** while the amount of TGFβ1 was higher in the supernatant from purified KRT17-eGFP^+^ than in that from KRT17-eGFP^-^cells **(Fig. S3I).** Both endogenous and exogenous TGFβ therefore induce KRT17-eGFP^+^ aBCs. Replating TGFβ1-induced KRT17-eGFP^+^ cells in iRAP conditions, with or without of TGFβi, reduced expression of KRT17-eGFP **(Fig. S3J)** and of aBC markers **(Fig. S3K)** while restoring expression of TRBC markers **(Fig. S3L)**. TGFβ therefore reversibly induces proliferation arrest and expression of canonical aBC markers in iRAPs.

At variance with our findings, human AT1 cells^55,84^ and aBCs^61^ were reported to derive from AT2 cells based on *in vitro* studies. We therefore examined the model systems used, hPSC-derived iAT2 cells and adult AT2 organoids, respectively. We procured human iAT2 cells, generated by repeated isolation and expansion of tdTomato^+^ SFTPC-reporter expressing cells in the same conditions as iRAPs.^85,86^ While expressing elevated levels of *SFTPC* mRNA, iAT2 cells expressed similar amounts of *SCGB3A2* mRNA **(Fig. S4A)** and protein **(Fig. S4B)** as iRAPs. When differentiated into iAT1 cells according to either our protocol (BMP4 and TGFβi)^62^ or to the protocol published by the same group (DCI and LATSi)^84^ the cells maintained elevated *SFTPC* **(Fig. S4C)** and tdTomato **(Fig. S4D)** expression. Similarly, TGFβ1 upregulated aBC genes in iAT2 cells, but only slightly downregulated TRBC genes in those compared to iRAPs **(Fig. S4E)**. TGFβ1 downregulated HT2-280 expression **(Fig. S4F,G)**, but did not downregulate TdTomato **(Fig. S4F,H)**. iAT2 cells are therefore more similar to iRAPs than to AT2 cells, but may be epigenetically dysregulated because of repeated reporter selection.

We next analyzed scRNAseq datasets from ferret distal lung organoids^87^ **(Fig. S4I)** generated using a protocol to expand human AT2 cells from adult lungs^55^ and from human distal lung organoids reported by Dost et al, 2026 **(Fig. S4J)**. While AT2 cells were present in both models, a large fraction of the cells expressed *SFTPB* together with SC and AT1 markers, suggesting that they are actually TRBC subsets. A determination of the role of AT2 cells in AT1 or aBC generation can therefore not be unequivocally made experimentally using these models, although computational analysis indicates that these trajectories exist **(Fig. 2)**.

### Cross-species comparison of aBCs, aBC-like cells and ADI cells

To gain insight into the molecular differences between TGF-β1 and LATSi-induced KRT17-eGFP^+^ cells, we performed hashed scRNAseq analysis **(Fig. 5A)**. TGFβ1-treated cells showed increased expression of aBC markers, while LATSi-treated cells expressed more AT1 markers. The TRBC markers, *SCGB3A2*, *SFTPB* and *SFTPC*, were reduced in both conditions **(Fig. 5B)**. These findings were consistent with RT-qPCR data **(Fig. 4F,G,H,I; Fig. S3G)**. Although the 200 genes most strongly induced by TGFβ1 induced genes included the ECM components, *FN1* and *COL1A1*, and the senescence marker, *CDKN2A*, regulation of neural projection, cell projection, cell migration and motility were the top-induced processes **(Fig. S5A)**. In LATSi-treated cells, on the other hand, tissue and epithelial development, response to hypoxia, wound healing and cell-substrate adhesion were among the top-induced processes, indicative of a regenerative response **(Fig. S5A)**. Live imaging confirmed increased motility of TGFβ1-induced KRT17-eGFP^+^ cells, while LATSi-induced cells were immotile **(Supplementary Movies 1-3)**.

**Figure 5.**
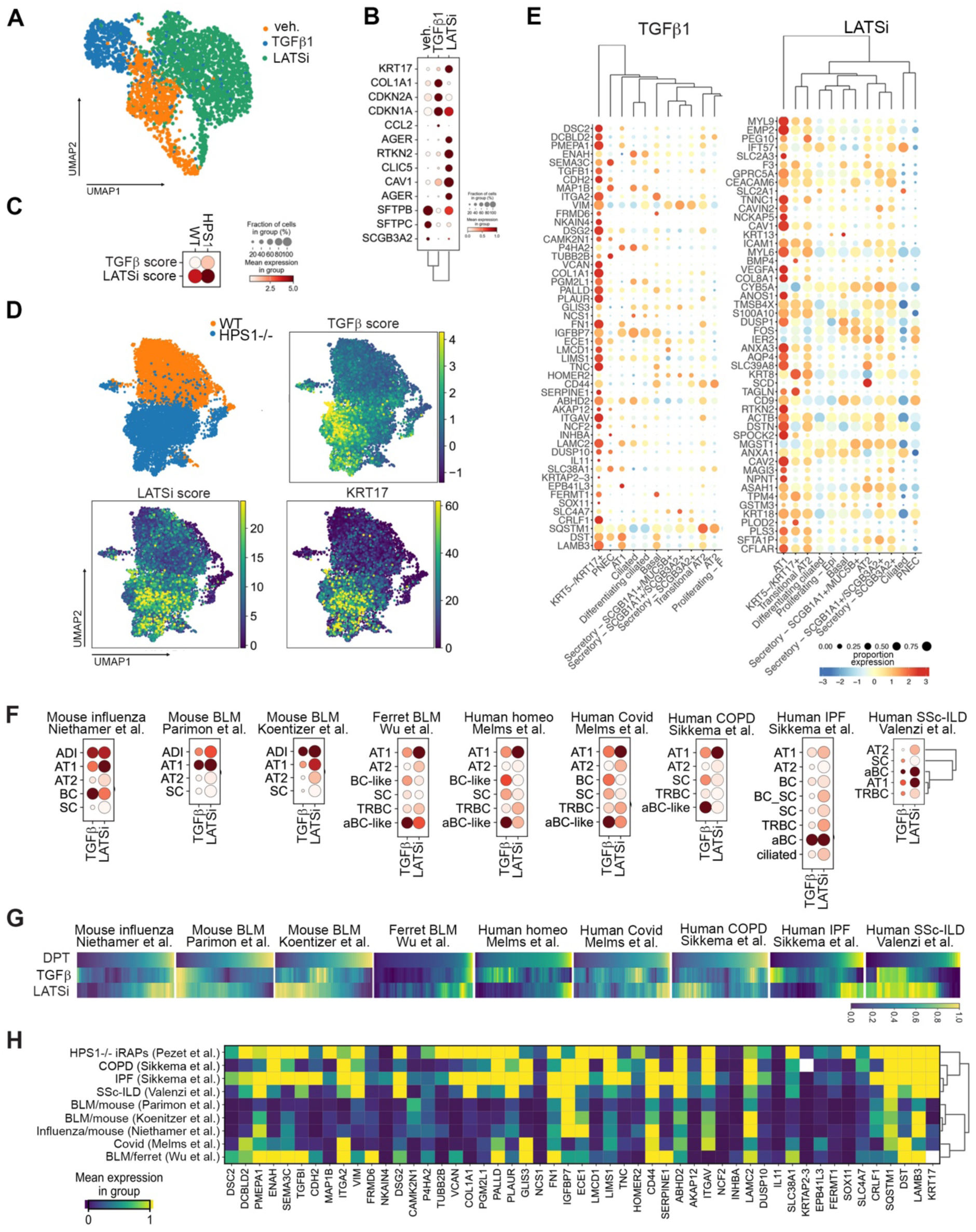
TGFβ and LATSi scores. **A.** UMAP of hash-tagged scRNAseq analysis vehicle, TGFβ or LATSi-treated iRAPs. **B.** Dot plot of TRBC, AT1 and aBC markers in vehicle, TGFβ or LATSi-treated iRAPs. **C.** Dot plot of TGFβ1 and LATSi scores in WT and *HPS1^-/-^*iRAPs. **D.** UMAP feature plots of integrated scRNAseq analysis of WT and *HPS1^-/-^*iRAPS for TGFβ and LATSi scores, and *KRT17*. **E.** Expression of TGFβ and LATSi signature genes in epithelial cells in LungMap IPF dataset. **F.** TGFβ and LATSi score in epithelial subsets in the datasets analyzed in Figs. 1 and 2. **G.** DPT analysis of TGFβ and LATSi signatures aBCs, ADI and aBC-like cells in the datasets analyzed in Figs. 1 and 2. **H.** Matrix plot of relative TGFβ signature gene expression in aBCs, aBC-like and ADI cells across datasets analyzed in Figs. 1 and 2.

We extracted TGFβ and LATSi gene signatures comprising the 50 top-upregulated genes in TGFβ1 and LATSi-treated iRAPs, respectively **(Fig. S5B),** to generate TGFβ and LATSi scores. We verified both scores in *HPS1^-/-^*iRAPs, where we previously showed accumulation of aBCs and a defect in AT1 differentiation.^62^ Both TGFβ and LATSi scores were upregulated in *HPS1^-/-^* compared to wild-type (WT) iRAPs **(Fig. 5C,D, Fig. S5C)**, suggesting that aBCs are characterized by elevation of both scores. To verify this finding *in vivo*, we examined expression of both signatures in the LungMap atlas.^88^ In normal lungs, the LATSi signature was associated with AT1 cells, as expected, whereas the TGFβ signature was not clearly associated with any specific population **(Fig. S5D)**. In the LungMap IPF dataset, on the other hand, the TGFβ signature was selectively and highly expressed in aBCs **(Fig. 5E)**. However, aBCs clustered next to AT1 cells with respect to expression of LATSi signature genes **(Fig. 5E)**. We then examined the TGFβ and LATSi scores in the lung injury datasets described in Fig. 2 **(Fig. 5F)**. Within each dataset, aBC-like cells and aBCs showed the highest TGFβ score, although only few cells expressed the TGFβ signature in the SSc-ILD dataset. The LATSi score was elevated in aBC-like cells in ferret BLM-treated lungs, and in human homeostatic and Covid lungs, but was the highest in AT1 cells. The similar pattern of TGFβ and LATSi scores in human and ferret aBC-like cells justifies their designation as such in ferrets, even though *KRT17* cannot be used as a defining marker in this species. In IPF and SSc-ILD, however, the LATSi score was the highest in aBCs. Mouse ADI cells also showed enrichment for both scores, although the TGFβ score appeared less specific for ADI cells in mice than for aBCs or aBC-like cells in humans **(Fig. 5F)**. Plotting TGFβ and LATSi scores in aBCs, aBC-like and ADI cells as a function of DPT showed out-of-phase waxing and waning of both scores **(Fig. 5G)**, explaining why the ADI, aBC-like and aBC clusters show enrichment for both scores. In SSc-ILD, however, this phenomenon was not observed and both scores were largely co-expressed **(Fig. 5G)**, likely indicative of the severe and pervasive AT1 differentiation defect in this condition. We conclude that, as a population, aBCs, aBC-like and ADI cells are defined by elevated TGFβ and LATSi scores, whereas at the level of individual cells, both scores are largely mutually exclusive.

Finally, as the TGFβ score was the most specific for aBCs, aBC-like and ADI cells, we compared these subsets with respect to this score, with inclusion of *HPS1^-/-^*iRAPs, by clustering based on identically scaled raw expression values of TGFβ signature genes **(Fig. 5H)**. aBCs in IPF, COPD and SSc-ILD and *HPS1^-/-^* iRAPs clustered together, further validating *HPS1^-/-^* iRAPs as an aBC model. aBC-like cells in Covid and in the ferret BLM model clustered with mouse ADI cells, indicating that they are distinct from aBCs in human fibrotic lung disease.

### Proximal airway cells generate injury-associated KRT17^+^ AT1 cells through an aBC-like stage

As PAGA analysis suggested input into the aBC-like subset not only from TRBCs, but also from BCs **(Fig. 2)**, we tested the hypothesis that human BCs possess AT1 potential through an aBC-like stage. Dissociation of LOs derived from ESCs or iPSCs and culture in conditions reported to expand epithelial stem cells from a variety of organs^89^ yielded continuously growing cell lines **(Fig. 6A,B)** with normal karyotype **(Fig. S6A,B)**. Comparison with the airway HLCA dataset ^67^ subsetted on epithelial cells revealed very close correspondence to the BC cluster (*KRT5^hi^TP63^hi^ITGB4^hi^*), as well as to cells transitioning between BCs and SCs (BC-SCs) **(Fig. 6C,D)**. We therefore called the cells ‘secretory and basal-like cells’ (SBLCs). BC-SCs in airways and in *in vitro* generated SBLCs expressed *KRT4* and *KRT13* and some *SCGB1A1*, the canonical SC marker **(Fig. 6D)**. *KRT4* and *KRT13* mark airway ‘hillocks’ in mice, considered intermediates between BCs and SCs^90^ and precursors of airway squamous metaplasia.^91^ Expression of BC markers (KRT5, KRT17 and TP63) was confirmed by IF **(Fig. 6E)**. Flow cytometry for CD104 (ITGB4) revealed CD104^hi^ (BC) and CD104^lo^ (BC-SC) populations **(Fig. 6F)**, consistent with scRNAseq analysis. SBLCs generated ciliated, goblet and secretory cells when cultured in air-liquid interface (ALI) conditions and therefore possess *bona fide* BC functionality *in vitro* **(Fig. S6C).** We next compared SBLCs with previously described hPSC-derived induced basal cells (iBCs) obtained by sequential selection using *NKX2.1* and *TP63* reporters.^92^ Integrated analysis showed that while *TP63* and *KRT17* expression was similar, *KRT5* expression was very low and *KRT4^+^* BC-SCs were absent in iBCs **(Fig. S6D,E)**. These findings indicate that SBLCs are more similar to canonical BCs and BC-SCs than iBCs are.

**Figure 6.**
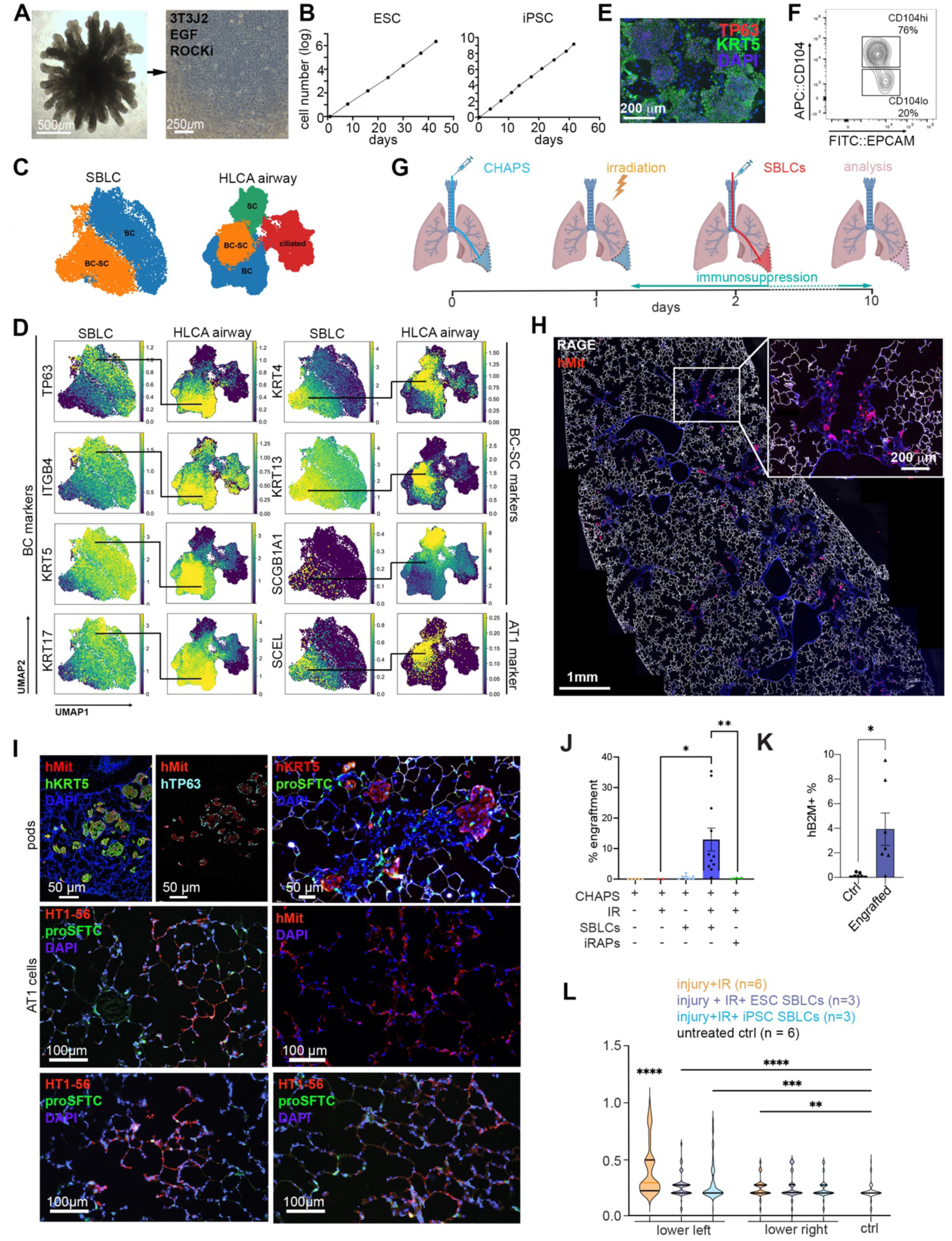
Generation and engraftment of SBLCs. **A.** Strategy for generation of SBLCs. **B.** Cellular expansion of ESC and iPSC-derived SBLCs. **C.** Cluster identity in SBLCs (integrated from ESCs and iPSCs) and in the airway subset of normal samples in HLCA. **D.** Feature plots of BC and SC markers in SBLCs and airway cells in HLCA. **E.** IF for BC markers. **F.** Flow cytometry of SBLCs for CD104. **G.** Engraftment strategy. **H.** Tile scan of engrafted lung 24h after transplantation. Arrows: hMit^+^ cells in injured (RAGE^-^) areas. **I.** IF staining at d10. **J.** Quantification of engraftment using quantitative human/rat DNA qPCR. n=4-11, Ctrl cells = LO cells, one-way ANOVA, **p < 0.01, ***p < 0.001. **K.** Quantification of engraftment by flow cytometry for hB2M on the dissociated lower left lung. n=6 (ESC and iPSC-derived cells), student’s t test; *, p<0.05; **, p<0.01. **L.** LIS analysis. IR: irradiation, UN: normal control lung, 800 blindly evaluated fields in total, 30 ROI per tile scan, one-way ANOVA; **, p<0.01; ***, p<0.001,****, p<0.0001.

Efforts to generate AT1 cells from SBLCs *in vitro* using our protocol for iRAPs^62^ or that of Burgess et al.^84^ for iAT2 cells failed. Attempts to engraft SBLCs in immune deficient *Nod.Scid.Il2rg^-/-^*(NSG) mice conditioned with polidocanol or naphthalene (airway injury) or BLM (alveolar injury), in some cases followed by irradiation to block endogenous repair, did not result in reproducible engraftment. Based on our previous study on *ex vivo* de-epithelialization of rat lungs^93^ we developed an *in vivo* model consisting of locoregional injury to the lower left rat lung using a mild detergent. scRNAseq showed striking accumulation of macrophage subsets **(Fig. S6F.i)**. After excluding hematopoietic cells **(Fig. S6F.ii)**, the relative proportions of all remaining cell types appeared preserved. Within the epithelial subset **(Fig. S6F.iii)**, ADI and proliferating AT2 cells appeared. Within the AT1 cluster, cells from control and injured rats showed only limited overlap **(Fig. S6F.iii)**, suggesting that AT1 cells in the injured rat were immature and most likely newly derived from ADI or from proliferating AT2 cells, near which they clustered. Furthermore, we observed a population of *Upk1a^+^* cells **(Fig. S6F.iii)**, which may correspond to variant secretory cells previously identified in the mouse.^33^ These data indicate selective epithelial injury accompanied by a vigorous inflammatory response together with AT2 proliferation and differentiation into AT1 cells.

We next used this injury model under immune suppression and after locoregional irradiation to eliminate the endogenous regenerative response **(Fig. S6G)** to engraft SBLCs by intrabronchial administration to the lower left lobe at d2 **(Fig. 6G)**. 24h after transplantation, SBLCs, identified by staining for human mitochondria (hMit), were present in injured areas characterized by the lack of RAGE staining **(Fig. 6H)**. At d10, patchy engraftment with human cells was observed **(Fig. S6H)**, with formation of hMit^+^hKRT5^+^hTP63^+^ pod-like structures similar to KRT5-pods observed in injured mouse and human lungs^38,40,94^ **(Fig. 6I)**. In addition, donor-derived alveolar ‘nets’ co-expressing HT1-56 were identified **(Fig. 6I)**. Fluorescence *in situ* hybridization did not identify cells containing both human and rat chromosomes, thus ruling out fusion **(Fig. S6I)**. Quantification at d10 by ratiometric qPCR for human Alu and rat DNA^95^ on consecutive sections **(Fig. S7A)** showed ∼5-10% engraftment **(Fig. 6J)**. iRAPs yielded ∼100-fold lower engraftment, possibly because of the absence of a TRB niche in rats, whereas engraftment was near-undetectable without irradiation **(Fig. 6J)**. Maximal engraftment sometimes occurred in the lower right lung **(Fig. S7A)**, likely due to spillover or anatomical variation in the airways, which could lead to inadvertent targeting of the right lung. Flow cytometry for hB2M on the lower left lung indicated a similar level of engraftment **(Fig. 6K)**. Blinded assessment of a standardized lung injury score (LIS)^96–98^ in a total of 800 randomly selected 1mm^2^ fields revealed that, whereas conditioned lungs remained injured at d10, administration of SBLCs resulted in a significant reduction in LIS **(Fig. 6L, Fig. S7B)**. Finally, transplantation of purified CD104^hi^ (BC) and CD104^lo^ (BC-SC) cells showed that all engraftment potential resided entirely in the CD104^hi^ BC fraction **(Fig. S7C-E)**.

Imaging-based spatial transcriptomics on the 10x Genomics Xenium platform, using a custom 100-target panel of probes each specific to either rat (which were also marked with mScarlet in this set of experiments) or human lung cells, revealed an engraftment pattern consistent with that seen by hMit staining **(Fig. S7F)**. mScarlet^+^ areas corresponded to either KRT5-pods or to alveoli **(Fig. 7A, top right; Fig. S7G)**. The pods were integrated into the surrounding host alveolar structure **(Fig. 7A, top right; Fig. S7G)** and contained consecutive layers of *hKRT5^+^* BC-like cells and *hSCEL^+^* AT1-like cells, with a fraction of the latter also expressing *hRTKN2*, towards the airspace **(Fig. 7A.1(a))**. *hKRT17* was expressed in *hKRT5^+^* BC-like cells **(Fig. 7A.1(b), Fig. S7G)**, but was also detected in some in airspace-facing AT1-like cells **(Fig. 7A.1(d-e))**. In addition, human *hSCEL^+^* and *hSCEL^+^hRTKN2^+^* AT1-like cells **(Fig. 7A.2, Fig. S7H)** were found integrated among rat AT1 cells (**Fig. 7A.1(d,e), Fig. 7A.2, Fig. S7H)**. *hAGER* was only detected very rarely **(Fig. 7A.2(f))**. All AT2 and AT1 cells were of rat origin, and these AT1 cells expressed *rAger* **(Fig. 7A, S7G,H)**. BCs therefore only possess AT1 potential, a finding consistent with trajectory analysis **(Fig. 2)**, while rat AT2 cells likely migrated to the injured areas, as has been previously shown in mice.^99^

**Figure 7.**
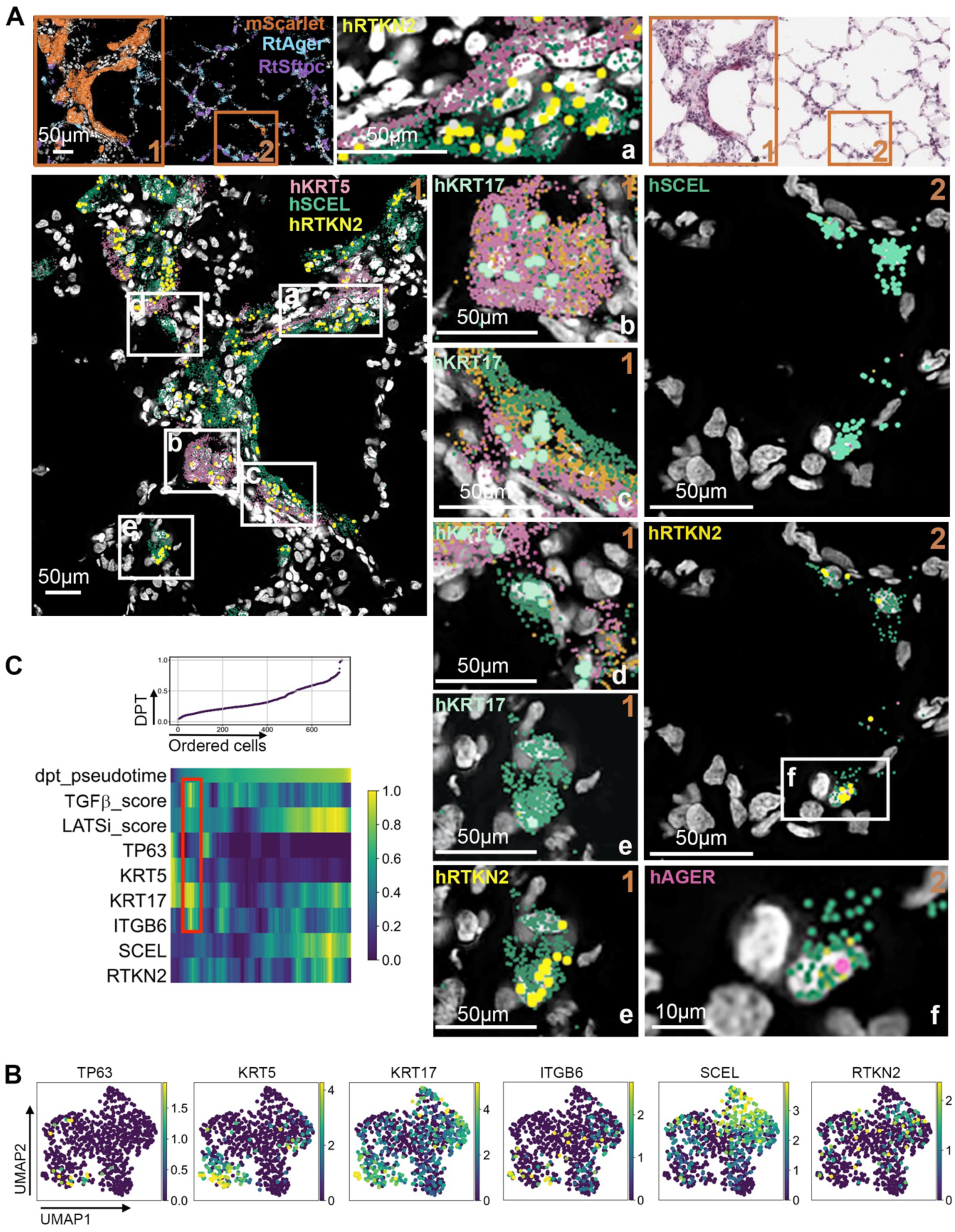
Transcriptomic analysis of engraftment. **A.** Representative example of Xenium spatial transcriptomics of an area containing a KRT5-pod differentiating into AT1-like cells **(1)** and human AT1-like cells integrated into rat alveolar structures **(2)**. Upper right: H&E of same area. **B.** Feature plots of scRNAseq analysis of mSCARLET^+^ engrafted human cells. **C.** DPT graph showing dynamic changes in expression of indicated markers as well LATSi and TGFβ scores over the DPT trajectory. Boxes: aBC-like stage.

scRNAseq on purified engrafted mScarlet-marked human cells showed a continuum of BC-like cells (*hKRT5^+^hTP63^+^*) to AT1-like *hSCEL^+^* and *hRTKN2^+^* cells, of which ∼40% expressed *KRT17* **(Fig. 7B).** Human AT2 cells were not identified. Plotting marker expression as a function of increasing DPT showed that the earliest cells in pseudotime expressed *hKRT5 and hTP63*, phenotypically corresponding to the engrafting BC-like cells, subsequently lose *hTP63*, *hKRT5* and *hKRT17*, but regain expression of *hKRT17*, before finally differentiating into *hKRT17^+^hSCEL^+^*AT1-like cells of which a fraction expressed *hRTKN2*^+^ **(Fig. 7C)**, a phenotype is similar to that observed in *hKRT17^+^* AT1 cells in Covid lungs **(Fig. 3M)**. Interestingly, cells at the intermediate *hKRT5^lo^hTP63^lo^hKRT17^+^*stage also expressed the aBC marker, *ITGB6*, and showed elevated TGFβ score and more mildly increased LATSi score **(Fig. 7C)**. This population is therefore consistent with an aBC-like state, supporting the prediction from computational analysis **(Fig. 2)** of a lineage relation between BCs and aBC-like cells.

Collectively, these findings indicate that BCs can give rise to aBC-like and AT1-like cells, consistent with the computational prediction.

## DISCUSSION

We demonstrate here the existence of regenerative trajectories in human and ferret lungs that are absent in mice and swine and may explain the challenge to recapitulate human distal lung diseases in mouse models.

Computational analysis indicates that AT1 regeneration from AT2 cells is conserved across all species studied, but is complemented in humans and ferrets by AT1 regeneration from TRBCs and from BCs. The inferred AT1 potential of human TRBCs and BCs was confirmed in organoid and xenotransplantation studies, respectively. Conservation of trajectories involved in lung regeneration therefore appears to align with the presence or absence of TRBs. Organoid studies have yielded conflicting results on the potential of TRBCs, however. Studies using adult organoids suggested that AT2 cells generate the most distal TRBC subset, known as AT0 cells, and then give rise to AT1 cells and to other TRBC subsets.^55^ Basil et al.,^53^ on the other hand, reported, based on studies in hPSC-derived airway organoids, the opposite trajectory, TRBCs>AT2 cells. A caveat in these studies is that iAT2 cells and adult lung-derived AT2 organoids appear more similar to TRBCs than to AT2 cells. We previously showed in iRAPs that TRBCs generate AT1 cells without AT2 intermediate.^62^ Our computational analysis suggests that each of these proposed trajectories exists *in vivo*. Furthermore, we previously demonstrated extensive plasticity among TRBC subsets present in iRAPs.^62^ Taken together, TRBC subsets and AT2 cells may in fact represent an extraordinarily plastic continuum of cell states capable of AT1 regeneration, a notion also supported by the striking transcriptomic coalescence of distal lung subsets after injury in human and ferrets.

While airway subsets participate in lung regeneration in both mice and humans, their nature and the trajectories engaged differ. Lineage tracing in mice consistently placed *Trp63^+^Krt5^+^* BC-like cells downstream of SCs or secretory-like subsets in trajectories towards AT2 cells,^34,36,39,44^ although a direct trajectory from SC-derived BC-like cells to AT1 cells was demonstrated in one report.^36^ In contrast, our data show that human BCs generate AT1 cells. The histological appearance of the structures formed by engrafting SBLCs resembles squamous metaplasia in the distal lung after acute lung injury in humans.^94^ suggesting that this lesion may at least in part represent emergency AT1 regeneration from migratory BCs.

Our study sheds light on the biology of human aBCs and aBC-like cells. The aBC-like phenotype represents a physiological, Hippo and TGFβ-regulated AT1 precursor that can be derived from AT2 cells, TRBCs and BCs and is, given its varied origins, best understood as a functional AT1-committed state. The aBC-like state is orthologous to mouse ADI cells which are, in contrast, primarily descended from AT2 cells, although derivation from airway SC has been reported^28^ as was also suggested by our PAGA analysis in the influenza model. aBCs, aBC-like and ADI cells transition between elevated LATSi or TGFβ signatures over pseudotime. As the TGFβ signature confers migratory capacity, while Hippo inhibition drives aBC-like cell differentiation from TRBCs, migration and differentiation may be temporally mutually exclusive. In line with this notion, terminal AT1 differentiation from iRAPs requires inhibition of endogenous TGFβ signaling.^62^ aBCs express the TGFβ signature genes more uniformly than aBC-like and ADI cells. As human and ferret aBC-like cells cluster with mouse ADI cells with respect to the TGFβ signature, the transient, inflammatory fibrotic response induced by BLM, classically used as a proxy for IPF, likely recapitulates features of human acute lung injury better than IPF.

TRBCs generate AT1 cells through two trajectories, inferred computationally in human datasets and verified functionally in iRAPs, that are distinguished by transit through an aBC-like stage and at least transient expression of KRT17. The apparent dominance of the KRT17^+^ trajectory in the iRAP model *in vitro* is likely a reflection of the fact that cytokine-supported, expanding organoids recapitulate a regenerative response rather than homeostasis. KRT17^+^ AT1 cells may differentiate into canonical KRT17^-^AT1 cells or may ultimately be replaced by AT1 cells derived from the KRT17^-^pathway. Consistent with the notion that the KRT17^+^ pathway is injury-associated, Hippo inhibition, which promotes AT1 regeneration in mice,^77,78^ induced KRT17^+^ aBC-like cells from iRAPs. This pathway may represent a more rapidly committing emergency AT1 regeneration mechanism, analogous to direct megakaryopoiesis, where megakaryocytes differentiate directly from hematopoietic stem cells to generate hyperreactive platelets in conditions of hematopoietic stress.^100–103^ An important implication of these findings is that therapeutic efforts to deplete aBCs in IPF should ideally preserve the function of physiological aBC-like cells.

Regenerative trajectories in human lungs are differentially affected by disease. Computational analysis suggests that TRBC>AT1 trajectories are predominant in Covid, either because of a more important role for TRBCs in distal lung regeneration, or, not mutually exclusively, because of impaired AT2>AT1 conversion. In contrast, the preservation of the AT2>AT1 trajectory together with disruption of TRBC>AT2 and TRBC>AT1 trajectories in IPF and, to a lesser extent, in COPD, indicates an important role for TRBC dysfunction in these diseases. While small airway disease is a well-established component of COPD^104^ and the presence of TRBCs likely explains why chronic bronchitis is recapitulated better in ferrets than in mice,^105,106^ TRBC dysfunction in IPF is more controversial. This notion is supported, however, by our observations that *HPS1^-/-^*iRAPs, which lack mature AT2 cells, show impaired AT1 differentiation^62^ and generate aBCs^62^ that, as we show here, cluster with aBCs with respect to the TGFβ signature. In addition, aBCs are directly connected to TRBCs and not to AT2 cells in PAGA analysis of IPF lungs, a finding consistent with the identification of SCGB3A2^+^SFTPB^+^KRT17^+^ cells positioned next to SCGB3A2^+^SFTPB^+^ TRBCs in IPF lungs.^62^ A role for defects in TRBCs in IPF pathogenesis is also supported by the observation that AT1 loss^59^ and morphological changes in small airways^107–109^ precede fibrosis. Furthermore, a gain-of-function promoter polymorphism in *MUC5B*, the most frequent, though lowly penetrant, genetic lesion in IPF,^110,111^ primarily leads to increased MUC5B expression in distal airways.^58,112,113^ Finally, occupational exposure to metal, stone and wood dust is a major IPF risk factor.^114^ These larger particles are deposited preferentially in airways.^115^ IPF pathogenesis may be heterogeneous as surfactant mutations may primarily affect AT2 cells.^8^ Pathogenic drivers of fibrotic lung disease may differentially impair AT1 regeneration from specific subsets within the TRBC-AT2 spectrum leading to accumulation of profibrotic aBCs. We do note, however, that iRAPs transiently co-express AT1 and AT2 markers with *SCGB3A2* (known as ‘transitional AT2’^54^ or ‘AT0’ cells^55^) in the canonical trajectory to iAT1 cells.^62^ TRBC function may therefore be affected by these mutations as well. Mutations in genes involved in telomere maintenance would impair the function of any regenerative subset. While expanding the regenerative arsenal of the distal lung, TRBCs may be a vulnerability in the context of fibrosis. This is suggested not only by the challenge to recapitulate IPF in mice, but also by the fact that spontaneous fibrosis is not known to occur in hooved domestic animals, which lack TRBs^51,53,55^ while cats and one dog breed^116^ which, like humans do have TRBs,^52^ develop an IPF-like disease (pet ferrets develop tumors and only live to 5-8 years). The conservation of regenerative trajectories and distal lung anatomy between ferrets and humans suggests that introduction of mutations associated with familial IPF in ferrets may yield more faithful animal models. In SSc-ILD, which is caused by dysregulated immunity,^117^ on the other hand, aBC-like>AT1, AT2>AT1 and TRBC>AT1 transitions were disrupted while TRBC>AT2, AT2>TRBC and BC>aBC-like trajectories were preserved, suggesting that the pathogenic process in SSc-ILD specifically impairs all terminal AT1 differentiation irrespective of the identity of the AT1 precursor. Inferred regenerative trajectories are therefore disrupted in a disease-specific fashion.

In conclusion, the more complex and only partially conserved regenerative trajectories in the human lung justify the use of hPSC-derived organoids and humanized animal models to dissect the unique features of normal and pathological human lung regeneration, and inform on the development of more appropriate animal models for human lung disease.

### Limitations

Although organoids are highly reductionist models, the concordance between *in vitro* studies, xenograft experiments and computational analysis supports the veracity of the unique regenerative mechanisms we identified in humans. Xenograft models are limited by the need for immune suppression, the species specificity of niches and signaling pathways, and often the shorter observation time. In the rat model described here, transplantation experiments were terminated at d10 because of weight loss and skin injury caused by the immune suppressive regimen containing tacrolimus. Slow-release pellets and osmotic pumps yielded erratic tacrolimus levels. An immunodeficient rat model that lacks T, B and NK cells^118^ showed more extensive lung injury after conditioning and very low engraftment. Lung regeneration may require signals originating from immune cells,^119–122^ possibly explaining the data. Despite these limitations, xenograft models have been essential for identifying human hematopoietic stem cells,^123^ recapitulating human immune responses,^124^ modeling human cancer^125,126^ and supporting the preclinical development of hESC-derived dopaminergic neurons for Parkinson’s disease.^127^

## Acknowledgments

This work was supported by the Kully Family Foundation (HWS) and grants NIH S10 OD032447 (HWS), Stony Wold-Herbert Fund (NVD, CP), and NIH T32GM145440 (JAT). ES received financial support from the Faculty of Medicine, Ruhr University Bochum, through the ICEP Scholarship 2025 for a one-year research stay at Columbia University, New York. This research was also funded in part through the NIH/NCI Cancer Center Support Grant P30CA013696 and used the Tissue bank part of the Molecular Pathology Shared Resource. The microscopy shared resource is partially supported by Cancer Center Support Grant P30CA016087. Professor Zähres is a supervisor of ES at the University of Bochum.

## Author Contributions

JAT performed all experiments in iRAPs, HYL generated and characterized SBLCs and analyzed engrafted lungs and spatial transcriptomics, CP developed the rat engraftment model, performed transplantations and analyzed engrafted lungs with HYL, ES worked with JAT on the iRAP model, ADS assisted JAT and ES, SL assisted with immunofluorescence on engrafted rat lungs, TAT assisted with bioinformatic analysis, JWM advised on microscopy, JW advised on spatial transcriptomics, YWC developed initial culture conditions and characterization of SBLCs, AS performed lung injury scores and pathology, NVD oversaw rat transplantations, HWS provided study concept, supervised experiments and data analysis, performed computational analysis with TAT, and wrote the manuscript.

## Data Availability

Newly generated data will be made available upon publication.

## METHODS

### Animals

Sprague-Dawley (SD) rats, 8-12 weeks old, males and females, weighing 180-250 g, were used in the experiments conducted to optimize and characterize the regional lung de-epithelialization and irradiation (n=36), and the transplant studies (n=14), respectively. All animal work was approved by the Columbia University Institutional Animal Care and Use Committee (IACUC), in compliance with the National Research Council Guide for the Care and Use of Laboratory Animals (eighth edition).

### Human lung samples

De-identified human lung samples were provided by the Herbert Irving Comprehensive Cancer Center Molecular Pathology Shared Resource Tissue Bank core (Institutional Review Board, protocol #AAAB2667). Normal controls consisted of pathologically normal tissue from patients undergoing procedures for pneumothorax. Acute lung injury samples were obtained from patients with a pathological diagnosis of diffuse alveolar damage.

### Human pluripotent stem cells (hPSCs)

Rockefeller University ES cell line 2 (female, passage 24–32) was maintained on mouse embryonic fibroblasts (MEFs) plated at 22,500 cells per cm^2^. Cells were cultured in human ES (hESC) cell maintenance medium consisting of DMEM/F12 (Thermo Fisher Scientific), 20% knockout serum (Stem Cell Technologies), 0.1 mM β-mercaptoethanol (Sigma-Aldrich), 0.2% Primocin (InvivoGen), 20 ng ml^−1^ *FGF2* (R&D Systems/Biotechne) and 1% Glutamax (Thermo Fisher Scientific), which was changed daily. Human ES cells were passaged every 3–4 days with Accutase (Innovative Cell Technologies) and were passaged at least two times before differentiation, washed and replated at a dilution of 1:24. Cultures were maintained in a humidified 5% CO2 atmosphere at 37 °C. Lines were karyotyped and tested for *Mycoplasma* contamination using PCR (InVivoGen) every 6 months.

#### Generation of lung organoids

Human LOs were generated as described previously. Briefly, MEFs were depleted by passaging 5–7 × 10^6^ ES cells onto Matrigel (Corning) coated 10-cm dish. Cells were maintained in hES cell medium in a humidified 5% CO2 atmosphere at 37 °C. After 24 h, cells were detached with 0.05% Trypsin/EDTA (Thermo Fisher Scientific) and distributed into six-well low-attachment plates containing primitive streak/embryoid body (EB) medium (10 μM Y-27632 (Tocris/Biotehne) and 3 ng ml^−1^ BMP4 (R&D/Biotechne)) to allow EB formation. EBs were fed every day with fresh endoderm induction medium (10 μM Y-27632, 0.5 ng ml^−1^ BMP4, 2.5 ng ml^−1^ *FGF2* and 100 ng ml^−1^ Activin A (R&D/Biotechne)) and maintained in a humidified 5% CO2/5% O2 atmosphere at 37 °C. Endoderm yield was determined by dissociating EBs and evaluating CXCR4 and c-KIT (Biolegend) co-expression by flow cytometry on day 4. Cells used in all experiments had an endoderm yield of >90% and were plated on 0.2% fibronectin-coated wells (R&D/Biotechne) at a density of 80,000 cells per cm^2^. Cells were incubated in anteriorization medium 1 (100 ng ml^−1^ Noggin (R&D/Biotechne) and 10 μM SB-431542 (Tocris)) for 24 h, followed by anteriorization medium 2 (10 μM SB-431542 (Tocris) and 1 μM IWP-2 (Tocris)) for another 24 h. At the end of anterior foregut endoderm induction, cells were switched to ventralization/branching medium (3 μM CHIR-99021 (Tocris), 10 ng ml^−1^ *FGF10* (R&D/Biotechne), 10 ng ml^−1^ recombinant human (rh)KGF (R&D/Biotechne), 10 ng ml^−1^ BMP4 and 50 nM all*-trans-*retinoic acid (Tocris)) for 48 h and 3D clump formation was observed. The adherent clumps were detached by gentle pipetting and transferred to a low-attachment plate, where they folded into lung bud organoids (LBOs) at days 10–12. The branching medium was changed every other day until days 20–25 and LBOs were embedded in 100% Matrigel in 24-well transwells (BD Falcon) inserts. The branching medium was added after the Matrigel solidified and changed every 2–3 days.

#### Generation of KRT17-eGFP reporter line

Alt-R™ technology (IDT) was used to design crRNAs and perform nucleofection. crRNA (100 μM) targeting the sequence 5’-GTCCACCAGACCACCCGCTG-3’ was combined at equal molar amounts (200 pmols) with Alt-R™ CRISPR-Cas9 tracrRNA-ATTO™ 550 (200 μM) and annealed at 95°C for 5 min in a thermocycler (Bio-Rad). The annealed duplex was then combined with Alt-R™ S.p. Cas9 Nuclease V3 (IDT) at a 3:1 molar ratio (duplex:Cas9) and incubated at room temperature for 10–20 min to form the ribonucleoprotein (RNP) complex. RUES2 cells (passage 22) were maintained on MEF-coated 6-wells plates until reaching ∼80% confluency. The cells were then collected with Accutase, washed, and resuspended in PBS. 1 x 10^6^ cells were transferred to a new tube, pelleted, and resuspended in 25 μL of P3 primary cell solution supplemented with supplement 1, as described by the manufacturer (Lonza, P3 Primary Cell 4D-Nucleofector Kit). To this, 1.25 μL of the RNP complex was added, along with Alt-R™ Cas9 Electroporation Enhancer at a final concentration of 1.75 μM (IDT), and 150 pmol of HDR donor Oligo. Cells were then transferred to cuvettes provided in P3 Primary Cell 4D-Nucleofector Kit, and nucleofection was performed using an Amaxa 4D-nucleofector with the hESC preset settings. Cells were replated onto MEF-coated 6-well plates in hES maintenance medium supplemented with 1.38 μM Alt-R™ HDR Enhancer V2 (IDT). After 12–24 h, medium was replaced with unmodified hES maintenance medium. After an additional 24 h, cells were collected with Accutase, stained for human CD90 (BioLegend), and single-cell sorted into MEF-coated 96-well plates (2–3 plates per sample) using a Sony MA900 cell sorter. Plates were centrifuged briefly at 100xg and returned to a normoxic incubator for 2 weeks for colony formation, with medium changed every other day. Clones were collected and genomic DNA was isolated for genotyping by PCR using the primers *KRT17* F1 InGene (5’-CCCCAGACTTACTCTTTATGCC-3’) and *KRT17* Insert R1 (5’-GGAGGATGGTAATATCTTGGGG-3’). PCR products were submitted to Genewiz for Sanger sequencing.

#### Generation of Induced Respiratory Airway progenitors (iRAPs)

Human lung organoids (LOs) were first generated as described in Matkovic Leko et al. (2023). Matrigel-embedded LOs were ready for iRAP generation on day 42 of development. The medium was removed from the transwell and 1 ml of 2 mg ml^−1^ dispase (Corning) was added to release LOs from the Matrigel for 30–45 min in a normoxic incubator. The LOs were transferred to a 15-ml conical tube and washed with wash medium to neutralize dispase and then centrifuged at 200xg for 5 min. The pellet was incubated with 1 ml of 0.05% trypsin–EDTA in a normoxic incubator for 10–12 min with occasional pipetting with a P1000 pipette tip. Single-cell dissociation was verified using a bright-field microscope. If a single-cell suspension was not obtained after 12 min, cells were washed with wash medium and incubated for an additional 5 min with 0.05% trypsin–EDTA. Cells were counted using a hemocytometer and 400 cells per μl of undiluted Matrigel were plated in a well of a 12-well non-tissue culture plate. The plate was placed in a normoxic incubator for 30 min until the Matrigel polymerized and 1 ml of CK-DCI medium (3 μM CHIR-99021, 10 ng ml^−1^ rhKGF, 50 ng ml^−1^ dexamethasone (Thermo Fisher Scientific), 0.1 mM 8-bromoadenosine 3′,5′-cyclic monophosphate sodium salt (Tocris) and 0.1 mM IBMX (Sigma-Aldrich)) was gently added using a 10-ml serological pipette. The medium was changed every 3 days. After 2–3 weeks, an iRAP culture was established that could be maintained for at least seven passages.

#### Generation of transitional cells

Seven to ten days after passaging, small iRAP spheres had formed. CKDCI medium was then replaced with DCISB medium (50 ng ml^−1^ dexamethasone (Thermo Fisher Scientific), 0.1 mM 8-bromoadenosine 3′,5′-cyclic monophosphate sodium salt (Tocris), 0.1 mM IBMX (Sigma-Aldrich) and 10 µM SB-431542 (Tocris)) for 10–14 days.

#### Generation of iAT1 cells

24-well tissue culture treated plates were coated with Matrigel at 1:30 dilution and left at 4 °C overnight. For the stepwise differentiation pathway, mature transitional AT2-like cells were used. For the direct differentiation pathway, mature iRAPs were used. In both cases, cells were released from Matrigel using dispase, washed and dissociated with 0.05% trypsin–EDTA, as described above. Matrigel was aspirated from the plate immediately before seeding the cells. Then, 2 × 10^5^ cells were resuspended in 0.5 ml of SB–BMP4 medium (10 µM SB-431542 (Tocris), 10 ng ml^−1^ BMP4 (Tocris) and 10 µM Y-27632 (Tocris)) and added into one well. The cells were cultured for 14-21 days in a normoxic incubator at 37 °C.

#### iRAP treatments (TGFβ, TGFβi, LATSi, TEADi)

For TGFβ1 treatment, recombinant human TGFβ1 (10 ng/ml; R&D Systems, Cat. No. 7754-BH) dissolved in 4 mM HCl was added to CKDCI medium. Cells were maintained under this condition for 5 days. Cells were then collected for analysis, sorted and analyzed, or sorted and replated in CK-DCI ± 10 μM SB-431542 for reversibility experiments. For LATSi treatment, TRULI (10 μM; MedChem Express, Cat. No. HY-138489) dissolved in DMSO was added to CKDCI medium and cells were maintained under these conditions for 7–10 days. Cells were then either passaged in CKDCI + LATSi to maintain treatment or passaged into CKDCI alone for reversibility experiments. For TEADi treatment, K-975 (10 μM; MedChem Express, Cat. No. HY-138565) dissolved in DMSO was added to CKDCI and maintained for 7–10 days before analysis. For TGFβi, SB-431542 (10 μM; Tocris, Cat. No. 16-141-0) dissolved in DMSO was added to CK-DCI and maintained for 10 days or added immediately upon replating of TGFβ1-treated sorted cells for reversibility experiments. Vehicle was DMSO at equal volumes as added to treatment conditions unless stated otherwise. For all conditions, treatment was initiated after sphere formation (7–10 days after passaging) and medium was refreshed every 2-3 days.

#### iAT2 cell culture

iAT2 cells were purchased from the Center for Regenerative Medicine (CREM) iPS core (Boston University, Boston, MA). Cells were received as mature organoids in suspension Matrigel in media. Upon receipt, Matrigel was pelleted by centrifugation (400g), digested with dispase, washed, and trypsinized as described above to make single cell suspensions. Cells were then seeded in undiluted Matrigel at 400 cells/µl on non-tissue culture treated 12 well plates and placed in a 37 °C incubator for 30 minutes to allow the Matrigel to solidify. CKDCI supplemented with 10 μM Y-27632 (Tocris) was added for the first 2 days after replating, followed by CKDCI alone, with the medium changed every 2-3 days. Stepwise differentiation into iAT1 cells was done as described above. Differentiation using the protocol Burgess et. al. (2024) was performed using 7 days of CKDCI followed by 12 days of DCI-LATSi. TGFβ treatment was done the same way as for iRAPs described above.

#### Generation of SBLCs

Matrigel-embedded lung organoids can be used to generate SBLCs when they reach day 42 (d)42 development. Lung organoids were released from Matrigel by incubating with dispase (1U/ml, Corning) for 30-45 minutes in a normoxic incubator. Organoids were transferred and washed in a 15 ml-conical tube with wash medium (IMDM, 5% FBS) to neutralize the protease, then centrifuged at 200 g for 5 minutes. The pellet was dissociated into small cell clump to single cell with 0.05% Trypsin/EDTA in a normoxic incubator for 10-12 minutes with occasional pipetting using a P1000 pipette. Dissociated cells were neutralized with wash medium, then centrifuged at 400 g for 4 minutes. Dissociated cells were seeded onto Mitomycin C-treated 3T3-J2 feeders (20,000 cells/cm^2^) and cultured with SBLC medium (DMEM: Ham’s-F12=2:1, 6% FBS, 250 ng/ml amphotericin B (Fisher Bioreagent, Pittsburgh, PA), 25 ng/ml hydrocortisone (STEMCELL Technology, Cambridge, MA), 5 µg/ml recombinant human insulin (PeproTech, Cranbury, NJ), 8 µg/ml cholera toxin (Sigma-Aldrich), and 5 µM Y-27632 Rock Inhibitor). SBLC medium was changed every other day. SBLCs were passaged every week and could be maintained for more than 20 passages with a normal karyotype.

#### Air-liquid interphase culture

10^6^ SBLCs were seeded onto 24-well trans-well insert and kept with SBLC media on the upper and lower chamber overnight. Cells were further cultured with PneumaCult™-Ex Plus Medium (Stem Cell Technologies) on the upper and lower chamber 48 hours, after which ALI was induced and cells maintained in PneumaCult™-ALI media (Stem Cell Technologies).

#### EdU incorporation

For *in vitro* studies, cells were incubated with 15 µM EdU (5-ethynyl-2′-deoxyuridine) for 20 h before fixation and permeabilization using reagents provided in the kit. Incorporated EdU was detected using the Alexa Fluor 647 picolyl azide provided in the kit and analyzed by flow cytometry.

#### TGFβ1 ELISA

The Invitrogen Human TGFβ1 ELISA kit (Catalog no. BMS249-4) was used. KRT17-eGFP- and KRT17-eGFP^+^ cells were sorted and replated at 400 cells/µl of Matrigel in a 25 µl droplet in a 24 well non-tissue culture-treated plate. After 7 days, the medium was conditioned by incubating the spheres with 400 µl CKDCI for 3 days. Conditioned medium was diluted 1:10 in Assay Buffer, acid-activated by adding of 20 μl 1N HCl with incubation for 1 hour at room temperature, followed by neutralization with 20 μl of 1N NaOH. The ELISA was then performed according to the manufacturer’s instructions using the provided microwell strips, biotin-conjugate, streptavidin-HRP, TMB substrate, and stop solution. Absorbance was measured at 450 nm as the primary wavelength and TGFβ1 concentrations were determined based on a 7-point standard curve.

#### RT–qPCR

Total RNA was extracted according to the manufacturer’s instructions using the Direct-zol RNA microprep kit (Zymo Research) and 500 ng of total RNA was reverse-transcribed using the qScript XLT complementary DNA (cDNA) SuperMix (Quantabio). Technical triplicates of a 15-μl reaction mix (for use in Applied Biosystems QuantStudio7 384-well System) were prepared with 3 µl of diluted cDNA and run for 40 cycles. Relative gene expression was calculated from the average cycle threshold (*Ct*) value, normalized to *GAPDH* as the internal control and reported as fold change (2^−ΔΔ*Ct*^).

#### Flow cytometry and cell sorting

iRAPs embedded in Matrigel were released by incubating the spheres with dispase for 30– 60 min, then washed and dissociated into single cells with 0.05% Trypsin/EDTA in a normoxic incubator for 10–12 min with occasional pipetting using a P1000 tip. The single-cell suspension was stained in polystyrene round-bottom 12 × 75-mm tubes (BD Falcon). Primary HT1-56 (Terrace Biotech; 1:200 dilution) and HT2-280 (Terrace Biotech; 1:200 dilution) antibodies were incubated with the cells for 1 h at room temperature. Cells were washed twice with fluorescence-activated cell sorting (FACS) buffer (PBS and 1% BSA) and centrifuged for 5 min at 420*g*. Fluorochrome-labeled secondary antibodies (Alexa Fluor 488 (A21042) or Alexa Fluor 594 (A21044) goat anti-mouse IgM, Thermo Fisher Scientific) were diluted in FACS buffer at 1:200 and added for 45 min in the dark. Cells were washed twice and resuspended in FACS buffer for flow cytometric analysis using Novocyte Quanteon (Agilent). Cell sorting was performed on a Sony MA900 (Sony).

For cells isolated from *in vivo* studies, perfused rat lungs were inflated and submerged in an enzyme cocktail (4mg/ml Pronase, 5U/ml Dispase, 1mg/ml Collagenase IV, and 1X Accutase, and 0.33U/ml DNaseI) and incubated at 37 °C for 30 minutes with gentle shaking. Tissues were further micro-dissected and incubated at 37 °C for 15 minutes. After centrifugation, the cell-tissue pellet was incubated with TrypLE at 37 °C for 20 minutes with vigorous pipetting. Single-cell suspension was filtered through 40-µm strainer and washed with DMEM/F-12 containing 10% FBS. Single cells were resuspended in FACS buffer for cytometric analysis on a Novocyte Quanteon (Agilent).

#### Histology and immunofluorescence (IF) staining

For analysis of cultured cells, a cell scraper was used to carefully detach the Matrigel droplet with embedded iRAPs from the bottom of the 12-well plate. The droplet was then transferred to an OCT histology mold and embedded in Tissue-Tek OCT (Sakura Finetek). Frozen samples were cut using a cryotome at the thickness of 10–12 μm, collected on adhesive microscope slides, air-dried, fixed in 4% paraformaldehyde and then washed twice for 5 min in 50 mM glycine to inactivate PFA, followed by washing in PBS. Samples were permeabilized for 10 min in 0.2% PBST (PBS + 0.2% Triton X-100 (Thermo Fisher Scientific), blocked by incubation in PBS containing 5% donkey serum and then incubated overnight with primary antibodies in 2% donkey serum. On the next day, samples were washed three times in PBS and 1% donkey serum and incubated with secondary antibodies for 1h at room temperature. Nuclei were stained with DAPI (Thermo Fisher Scientific). Sections were mounted with Mounting Reagent (Agilent) on coverslips. Samples were imaged using a Leica TCS SP8 Stellaris Laser scanning confocal microscope and Leica DMi8 inverted phase-contrast microscope (Leica Microsystems). For IF signal intensity analysis, Fiji (ImageJ 1.54p) was used. For each analysis, 4 regions of interest (ROIs) from at least three biological replicates were analyzed, with at least 100 cells per ROI. Thresholds were applied uniformly across all analyzed images. Integrated density for each channel of interest was divided by the number of cells, which was determined using a DAPI-based mask. For YAP+ nuclei analysis, the DAPI mask was applied to the YAP channel and only YAP staining that overlapped with DAPI staining was counted.

For the rat studies, frozen sections (5µm thick) and H&E stainings from each experiment were prepared by the Molecular Pathology Shared Resource (MPSR) facility at Herbert Irving Comprehensive Cancer Center (HICCC),Columbia University, New York. Upon returning to the room temperature, the sections were submerged in 4% paraformaldehyde for 10 minutes, washed with PBS for 5 minutes, and permeabilized in 0.25% Triton X-100/PBS solution for 20 min. After one hour of blocking in 10% donkey serum/PBS solution, EdU staining was performed in the experiments where *in vivo* labeling was performed. Subsequently, primary antibodies were added to the sections diluted in 5% donkey serum/PBS solution. The slides were then incubated overnight in the dark at 4°C. The next day, following three 10-minute washes in 0.025% Triton X (ThermoFisher) -100/PBS solution, the sections were treated with secondary antibodies (in 5% donkey serum/PBS) for a 1hr in the dark. Slides were again washed three times for 10mins in 0.025% Triton X-100/PBS solution. DAPI staining (1:1000 in PBS) was performed for 10mins and the slides washes for 5mins in PBS before being mounted and sealed with nail polish. IF images were taken and processed using a Leica DMi8 system and with the Leica Stellaris for confocal imaging.

#### Live Imaging

iRAPs were plated in Matrigel domes at 400 cells/µl onto 12 well non-tissue culture-treated plates and maintained for 15 days in CKDCI. Cells were then switched to medium containing CKDCI+vehicle, CKDCI+LATSi, or CKDCI+TGFβ1 as described above. Cells were returned to the 37°C normoxic incubator conditions for 24 hours. Before live imaging on the Axio Observer Z1 (Zeiss): the imaging station was warmed to 37 °C and set to 5% CO2. Plates were then imaged individually. Images were taken using the 10x objective every 30 minutes for 96 hours. Images were processed in ImageJ (Fiji), removing out of focus or overexposed frames, and exported as .AVI files.

#### RNA-seq and data analysis

Total RNA was extracted from sorted KRT17-eGFP-and KRT17-eGFP^+^ CKDCI iRAPs using the Direct-zol RNA microprep kit Clontech Ultra Low v4 kit was used for cDNA synthesis followed by NexteraXT. Libraries were then sequenced using Element AVITI at Columbia Genome Center. The samples were multiplexed in each lane, which yield targeted number of paired-end 75bp reads reads for each sample. bases2fastq version 2.2.0.1940753917 was used for converting BCL to fastq format, coupled with adaptor trimming. Pseudoalignment was done to a kallisto index created from transcriptomes containing coding sequences (cDNA) (Ensembl v113, Human:GRCh38.p14). Differential gene expression was analysed using DESeq2 version 1.42.1 R package. DEGs were identified for our studyby setting a cutoff of *P* value < 1 × 10^−10^ and and fold change > ±2. GO analysis was performed using the DEGs with clusterProfiler package version 4.10.1, using genome wide annotation for human database (org.Hs.eg.db) version 3.18.0.

#### scRNAseq sample preparation

Direct and Stepwise scRNA sequencing: iAT1 cells from both pathways were prepared as described above and cultured in the same iAT1 condition medium for 2 weeks. Cells were then treated with trypsin for 8 minutes, collected and resuspended in 0.1%BSA. Cells were then stained with TotalSeq-B anti-human hashtags 1 and 2 for 3 minutes on a rotator. Cells were then washed 5 times in PBS and equal numbers of cells were combined. The viability of single cells was assessed using Trypan blue staining and debris-free suspensions with >80% viability were deemed suitable for scRNA-seq. Cells were then submitted for scRNA sequencing.

TGFβ and LATSi treated iRAPs: Samples were cultured in CKDCI for 2 weeks before being treated with DMSO, TGFβ, or LATSi at the concentrations described above. Samples were then released from Matrigel using dispase and dissociated into single cells using trypsin for 10 minutes. Samples were then multiplexed using the same protocol described above.

Single-cell gene expression profiles for ESC-derived and iPSC-derived SBLCs were generated with the 10x Genomics Chromium Single-Cell 3′ RNAseq platform, using an input of ∼10,000-20,000 cells. The Columbia University Genomic core performed raw data processing with 10x Genomics Cell Ranger pipeline. Briefly, gel bead-in emulsions (GEMs) were generated on the sample chip in the Chromium controller. Barcoded cDNA was extracted from the GEMs by post-GEM RT cleanup and amplified for 12 cycles. Amplified cDNA was fragmented and subjected to end repair, poly(A) tailing, adaptor ligation and 10x-specific sample indexing following the manufacturer’s protocol. Libraries were quantified using Bioanalyzer (Agilent) and QuBit (Thermo Fisher Scientific) analysis and were sequenced in paired-end mode on a NovaSeq instrument (Illumina) targeting a depth of 50,000–100,000 reads per cell.

#### scRNA-seq computational analysis

Sequencing data were aligned and quantified using the Cell Ranger single-cell software suite (version 6.1.2, 10x Genomics) against the provided GRCh38 (Ensembl 98) human reference genome. Publicly available datasets were downloaded from the GEO database or from the HLCA database **(Supplementary Table 1)**. Data were analyzed using Scanpy 1.11.5, SCVI 1.4.0, Pandas 2.3.3, Seaborn 0.13.2, and Numpy 2.3.5. Data were loaded into AnnData objects and filtered based on the minimum number of cells in which a gene was expressed, and the minimum number of genes per cell. Doublet exclusion was performed using SCVI. Cells were next filtered based on the percent of ribosomal and mitochondrial reads, and outliers in the number of genes per cell were excluded at the 98^th^ percentile. Individual samples were then concatenated and integrated using SCVI. which yields SCVI-normalized count data. The HLCA datasets were already integrated. Neighbors, UMAP, and Leiden clustering were next calculated using the X-scVI layer. Data were then subsetted on epithelial cells expressing canonical lung markers and not expressing markers for mesenchymal, hematopoietic and vascular cells. After subsetting neighbors, UMAP, and Leiden clusters were each time recalculated. The map function implemented in Scanpy was used to assign cell names to clusters using a mapping dictionary. Cell identity was assigned based on canonical markers. Next, PAGA analysis as implemented in Scanpy was performed with a resolution of 0.03 throughout. Data were subsequently reclustered using PAGA initialization. For diffusion pseudotime plotting, a root cluster was set, and a root cell was randomly chosen.

#### Xenium spatial transcriptomics

Spatial transcriptomic profiling was performed on frozen lung tissue from rat xenotransplantation experiments using the Xenium In Situ platform (10x Genomics). Engrafted rat lungs were embedded in optimal cutting temperature (OCT) compound, and 10-μm-thick sections were prepared and mounted on Xenium-compatible slides. Spatial gene expression was profiled using a custom, 100-gene YEFCKN species-specific Xenium panel (10x Genomics) containing probes targeting either human or rat transcripts, enabling identification and spatial characterization of human engrafted cells within the rat lung tissue.

Xenium processing, transcript detection, cell segmentation, and spatial localization of transcripts were performed according to the manufacturer’s protocol. Quality control and exclusion of low-quality cells were performed based on transcript detection and cellular segmentation metrics generated by the Xenium analysis pipeline. Spatial expression patterns and cellular distributions were subsequently visualized in Xenium Explorer (10x Genomics).

### Animal Studies

#### Anesthesia and euthanasia

Rats, both males and females, were anesthetized with isoflurane vapor (3-5%) and ketamine (VEDCO, Saint Joseph, MO) (80-95 mg/kg) and xylazine (Covetrus, Portland, ME) (5-10 mg/kg) by intraperitoneal (IP) injection. The animals were then positioned upright on a rodent workstand (Hallowell EMC, Pittsfield, MA) and endotracheally intubated with a modified cannula (JorVet, Loveland, CO). The cannula had a tightly fitting tracheal plug (cut from a 200 mL pipette tip) to seal the trachea during removal of the de-epithelialization solution. Throughout the experiment, animals, maintained on 1% isoflurane, were spontaneously breathing through the lung (right) that was not intubated and mechanically ventilated through the intubated lung (left). The left lung was ventilated using synchronized intermittent mandatory ventilation (SIMV) on a volume control modality for single lung ventilation (Tidal volume 3cc/Kg; PEEP 5; PIP limit 35) using the Inspira Asv Advanced Safety Animal Ventilator (Harvard Apparatus, Boston, MA). Euthanasia was performed with isoflurane (Covetrus) (5%, 15min) inhalation, followed by bilateral thoracotomy. Following a transverse incision of the aorta to facilitate blood release from the left ventricle, the main pulmonary artery was perfused with phosphate-buffered saline (PBS, Sigma-Aldrich, St. Louis, MO) via the right ventricle with a static pressure that was maintained at 13 cm H2O above the heart until the perfused lungs were blanched. OCT compound (Sakura Finetek, Torrance, CA) was diluted in PBS (8 OCT:2 PBS by volume) and injected into the trachea. The injection volume was equivalent to 60% of the animal’s total lung capacity (TLC, see below for calculation). A total of ten tissue samples were taken from each animal and embedded in the undiluted OCT compound for frozen sections.

#### Calculation of Total Lung Capacity (TLC) and relative de-epithelialization solution dose

We adopted a published tissue-volume equation of fixed lungs to calculate TLC in live SD rats ^128–130^. To apply the de-epithelization of up to 20% of the left lobe, the volume of the CHAPS-solution was calculated according to the formula optimized to cover 17-20% of the alveolar surface of the total left lung (derivation not shown): Dose (µL) = 6.3*W^0.7^ (W, body weight in g).

### *In vivo* de-epithelialization

The de-epithelialization solution contained 4 mM 3-[(3-cholamidopropyl)dimethylammonio]-1-propanesulfonate (CHAPS, Sigma-Aldrich), 0.5 M NaCl (Sigma-Aldrich), and 25 mM EDTA (Gibco, Waltham, MA) in deionized water. Each dose was instilled in two equal aliquots, 30 minutes apart. 30 minutes after the second instillation, the workstand was lowered to its horizontal plane position. After 15 minutes, bronchoalveolar lavage (BAL) was performed twice (1 mL of PBS each) via a 3-mL syringe. The tracheal plug was pushed down the cannula to create a temporary seal the tracheal entrance to facilitate the BAL. The plug was quickly retrieved after each lavage to ensure ventilation. The rats were kept on the ventilator for an additional 15-30 min while anesthesia was weaning, then extubated, and returned to their cages.

#### Monitoring and blood analysis

Heart rate, oxygen saturation, and rectal temperature were monitored during each experiment using a pulse oximeter (8500, Nonin Medical, Plymouth, MN) and a digital thermometer, respectively. Tail vein blood samples were collected one day before and 48 hours after de-epithelialization experiments and analyzed by the Columbia University Comparative Pathology Laboratory (CPL). In the engraftment experiments, the plasma level of tacrolimus was measured in blood samples collected from experimental and control rats the day of cells delivery and at 7 days post-transplant. Plasma was processed with the Abbott specific kit for tacrolimus level, according to the kit manufacturer’s protocol (Abbott, Abbott Park, IL) and analyzed using the Abbott Architect device. All samples with a tacrolimus level above 15-20 ng/ml were considered optimal for rat immunosuppression.

#### Lung irradiation

One day after de-epithelialization, rats were anesthetized with isoflurane (1-3%) and placed on the animal holder of the Small Animal Radiation Research Platform (SARRP). After a total body computerized tomography (CT) scan, the lung area was contoured and selected for irradiation using build-in software on the platform. Each rat received 12Gy delivered into two beams, one directed from above and the other from below the animal.

#### Cell transplantation

For the transplantation studies, rats were started on a daily triple drug immunosuppression regimen consisting of with Tacrolimus (4mg/kg daily subcutaneous, Prograf from Astellas, Northbrook, IL), Mycophenolate (0.1 mg/500 ml water, oral, Accord, Durham, NC), and Methylprednisolone (20 mg/500 ml of water, oral, Sagent, Schaumburg, IL) 24 hours after de-epithelialization and 24 hours prior to administration of the cells. 6 hours prior to cell delivery, rats were also treated with Methylprednisolone (5mg/kg, intramuscular). Rats were anesthetized and intubated with the same modified cannula used for the regional de-epithelialization (JorVet) targeting the lower left lobe using a flexible bronchoscope (1800Endoscope, Bradenton, Florida). 10^7^ SBLCs resuspended in culture media were delivered through the cannula divided in two boluses, with the animal ventilated for 15 min between them. Rats were weaned from anesthesia, returned to their cages, and euthanized as described above 10 days after the de-epithelialization.

#### Lung Injury Score (LIS)

Hematoxylin and Eosin (H&E) staining was performed by the Molecular Pathology Core (MP/MPSR, Columbia University Medical Center) using a standard protocol. The slides were scanned using a bright field whole slide scanning system (Leica AT2 digital scanning system) with a pixel size of 0.25µm. We then developed and applied a custom script (available here, https://codebeautify.org/alleditor/y234a8635) to select random and non-overlapping regions of interest from whole slide image scans of lung sections. A section from the targeted left lobe and a section from the contralateral right lobe (control) were used for the analysis. The script was used to obtain 30 regions of interest (ROI) per sample. Each ROI was then blindly evaluated for LIS according to the guidelines of the 2011 ATS Workshop Report on Features and Measurements of Experimental Acute Lung Injury in Animals ^96^ by a lung pathologist (AS).

#### Genomic qPCR

Genomic DNA was extracted from 200-250µm-thick cryopreserved rat lung tissue using the Zymo Quick-DNA MicroPrep Kit (Zymo, Irvine, CA), according to the manufacturer’s instructions. Tissue samples were washed with water to remove residual OCT before lysis with Zymo Genomic Lysis Buffer. DNA concentration was assessed by absorbance in a spectrophotometer (Thermo Scientific NanoDrop2000c). Human and rat DNA were verified by qPCR using human AluYb8 primers (*Alu,* Fw 5’-CGAGGCGGGTGGATCATGAGGT-3’, Rv 5’-TCTGTCGCCCAAGGCCGGAGCT-3’) and rat *aryl hydrocarbon receptor* gene (*Ahr* Fw 5’-TGCCCAGCAACAGCCTGTGAG-3’, Rv 5’-AACTGGCGAACATGCCATTGA-3’) ^131^ Serially diluted human genomic DNA (0.01pg-10ng) was mixed with 100ng of rat genomic DNA to determine a standard curve at each set of qPCR primers. PCR was performed on QuantStudio 5 Real-Time *PCR* System. qPCR was performed with denaturation at 95°C for 10 minutes, followed by 40 cycles of amplification of 95°C for 15 seconds, 62°C for 5 seconds and 72°C for 15 seconds. To quantify human cell engraftment in rat lungs, human AluYb8 qPCR was performed in 100ng of gDNA extracted from each section of the rat lung. Human gDNA quantity was calculated from raw Cq value based on the standard curve generated for each batch. The percentage of human cell engraftment was calculated as the amount of human gDNA relative to the total 100ng of gDNA extracter per rat tissue section.

### Quantification and Statistical analysis

Statistical analysis was performed using an unpaired two-tailed, Student’s t-test, one-way analysis of variance (ANOVA) or two-way ANOVA with multiple-testing correction where appropriate using Prism 9. Results are shown as the mean ± SEM. P values < 0.05 were considered statistically significant. The n value refers to biologically independent replicates. The statistical details of individual experiments are described in the Figure Legends.

## Legends to Supplementary data

**Figure S1.**
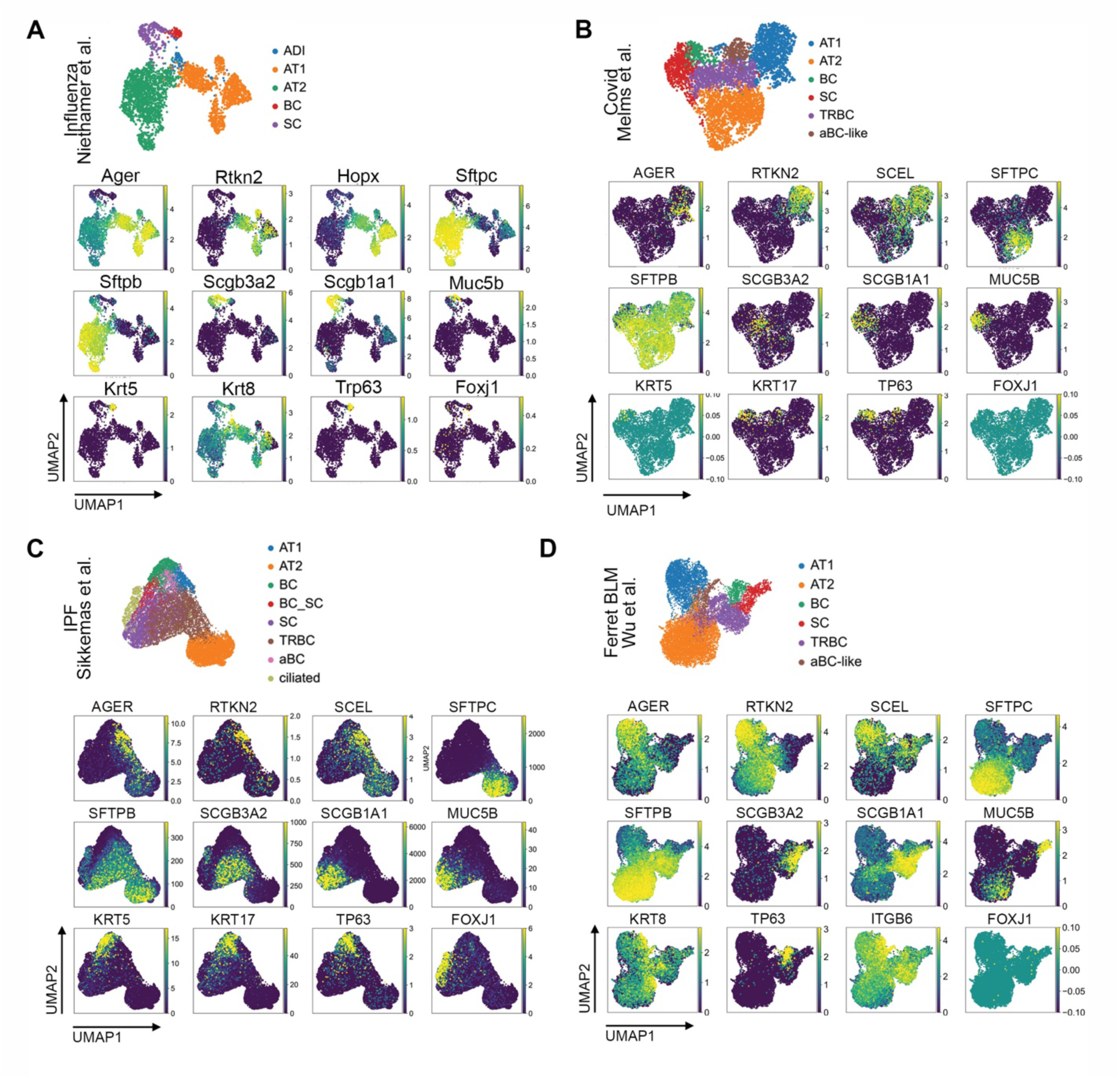
Cell type assignment in single cell transcriptomic datasets. Representative UMAP feature plots for lineage-defining markers in mouse influenza (**A**), Covid (**B**), IPF (HLCA, Sikkema et al., 2023) (**C**) and ferret BLM **(D)** datasets.

**Figure S2.**
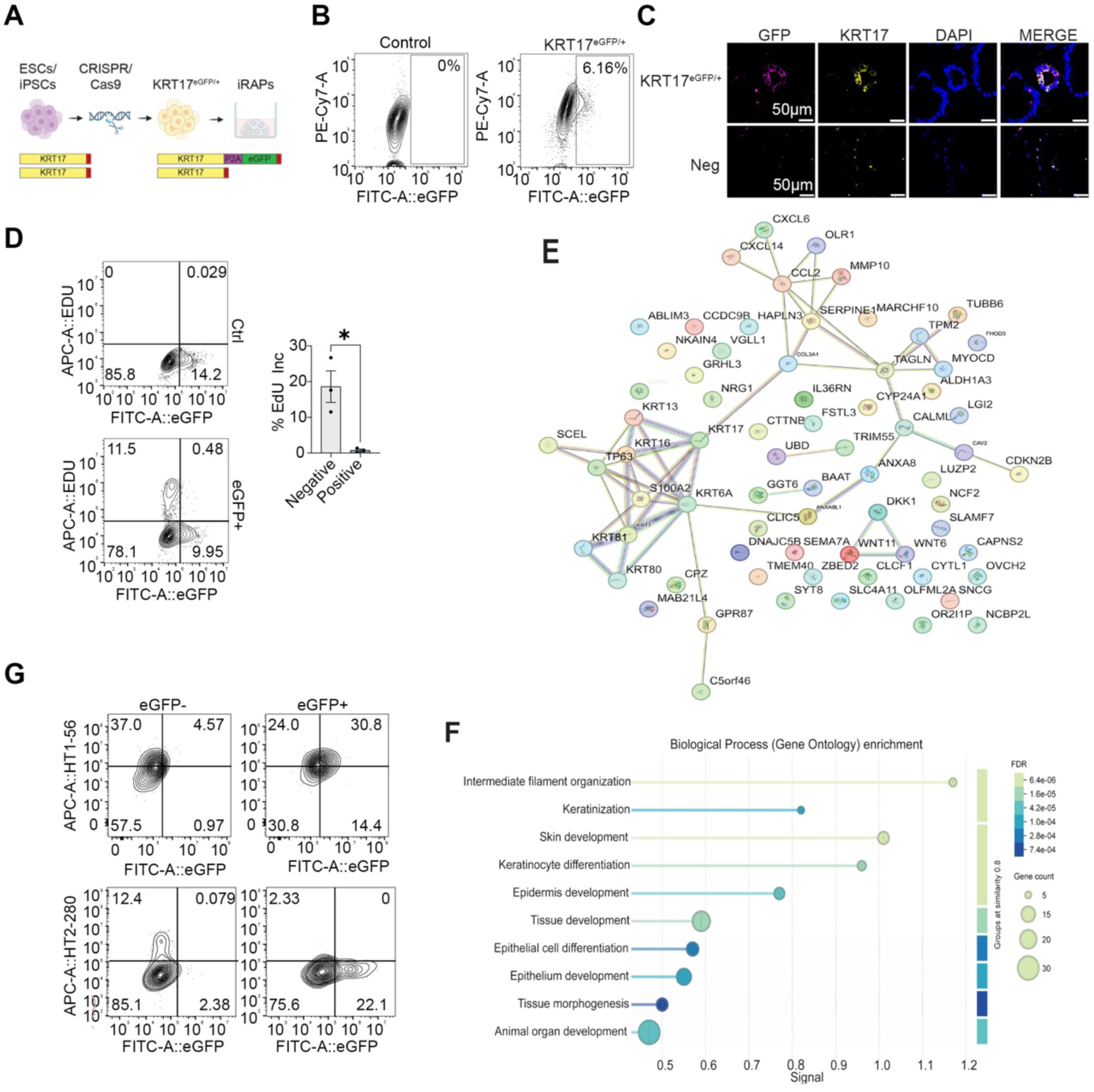
KRT17 reporter and identification of a KRT17^+^ AT1 lineage. **A.** Schematic of generation of KRT17-eGFP reporter. **B.** Representative flow plot of eGFP expression in iRAPs derived from KRT17-eGFP reporter hESC line. **C.** IF representative of 6 biological replicates of KRT17-eGFP reporter iRAP (top) and control fibroblasts (bottom) showing colocalization of labelled markers. **D.** Representative flow cytometry plots (left) and quantification of EdU^+^ cells (right). *p < 0.05, unpaired two-sided Student’s t-test, n=3. **E.** STRINGDB interaction network of the 71 differentially expressed genes between KRT17-eGFP^-^and KRT17-eGFP^+^ iRAPs as measured by bulk RNAseq (n=3). **F.** Gene ontology pathway analysis of upregulated genes in KRT17-eGFP^+^ iRAPs. **G.** Representative flow cytometry plot of expression of HT1-56 and HT2-280 in transitional cells generated from KRT17-eGFP^-^and KRT17-eGFP^+^ iRAPs.

**Figure S3.**
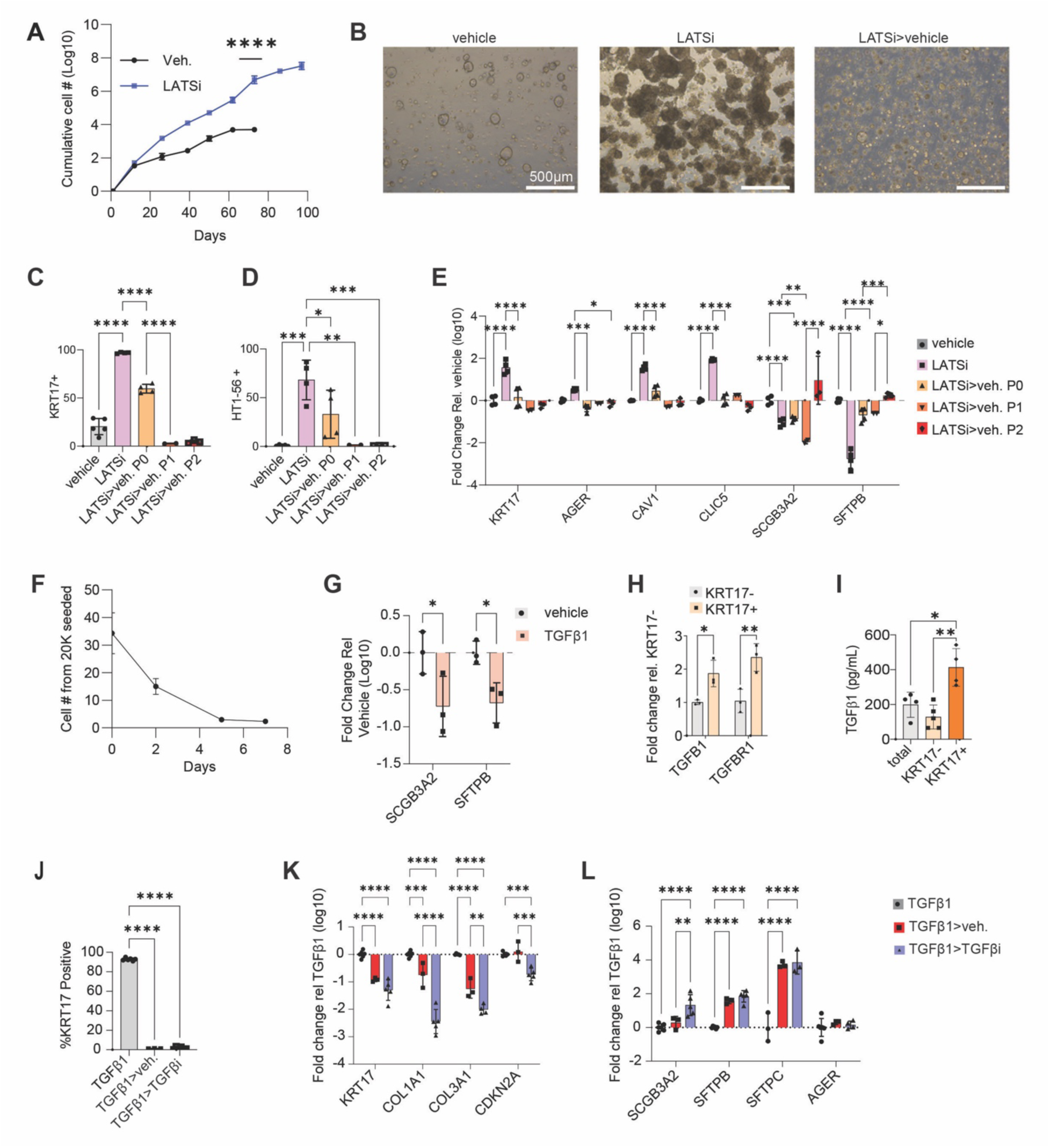
Reversibility of LATSi and TGFβ effects in iRAPs. **A.** Cumulative cell number in vehicle and LATSi treated iRAPs over 10 passages. ****p<0.0001, unpaired two-sided Student’s t-test, n=3. **B.** Representative brightfield images of iRAPs in vehicle, LATSi and LATSi>vehicle reversal conditions. **C.** Fraction KRT17-eGFP^+^ cells in LATSi and reversal conditions. ****p<0.0001, one-way ANOVA with Tukey’s multiple comparison test, n=5 (vehicle), n=4 (LATSi and LATSi>Veh. P0), n=2 (LATSi>veh. P1), and n=3 (LATSi>veh. P2). **D.** Fraction HT1-56^+^ cells in LATSi and reversal conditions. *p<0.05, **p<0.01, ***p<0.001, one-way ANOVA with Tukey’s multiple comparison test, n=5 (vehicle), n=4 (LATSi and LATSi>veh. P0), n=2 (LATSi>veh. P1), and n=3 (LATSi>veh. P2). **E.** mRNA expression of AT1 and TRBC markers in LATSi and reversal conditions. *p<0.05, **p<0.01, ***p<0.001, ****p<0.0001, two-way ANOVA with Tukey’s multiple comparison test, n=4 (vehicle), n=4 (LATSi and LATSi>veh. P0), n=2 (LATSi>veh. P1), and n=3 (LATSi>veh. P2). **F.** Cell number in TGFβ1-treated iRAPs over 7 days, n=3. **G.** mRNA expression of TRBC markers in TGFβ1 and vehicle-treated iRAPs. *p<0.05, two-way ANOVA with Šidák multiple comparison test, n=3. **H.** *TGFB1* and *TGFBR1* mRNA expression in sorted KRT17-eGFP^-^and KRT17-eGFP^+^ cells. *p<0.05, **p<0.01, two-way ANOVA with Šidák multiple comparison test, n=3. **I.** TGFβ1 protein concentration in conditioned media from sorted and replated total, KRT17-eGFP^-^ and KRT17-eGFP^+^ iRAPs. *p<0.05, **p<0.01, one-way ANOVA with Tukey’s multiple comparison test, n=4 (total and *KRT17*^+^) and n=5 (*KRT17*^-^). **J.** Fraction KRT17-eGFP^+^ cells in TGFβ1-treated cells after sorting and replating in vehicle or TGFβi conditions. ****p<0.0001, one-way ANOVA with Dunnett’s multiple comparison test, n=6. **K.** mRNA expression of aBC markers in TGFβ1-treated, and TGFβ1-treated, sorted and replated conditions. **p<0.01, ***p<0.001, ****p<0.0001, two-way ANOVA with Tukey’s multiple comparison test, n=3. **L.** mRNA expression of TRBC and AT1 markers in TGFβ1-treated, and TGFβ1-treated, sorted and replated conditions. **p<0.01, ****p<0.0001, two-way ANOVA with Tukey’s multiple comparison test, n=3.

**Figure S4.**
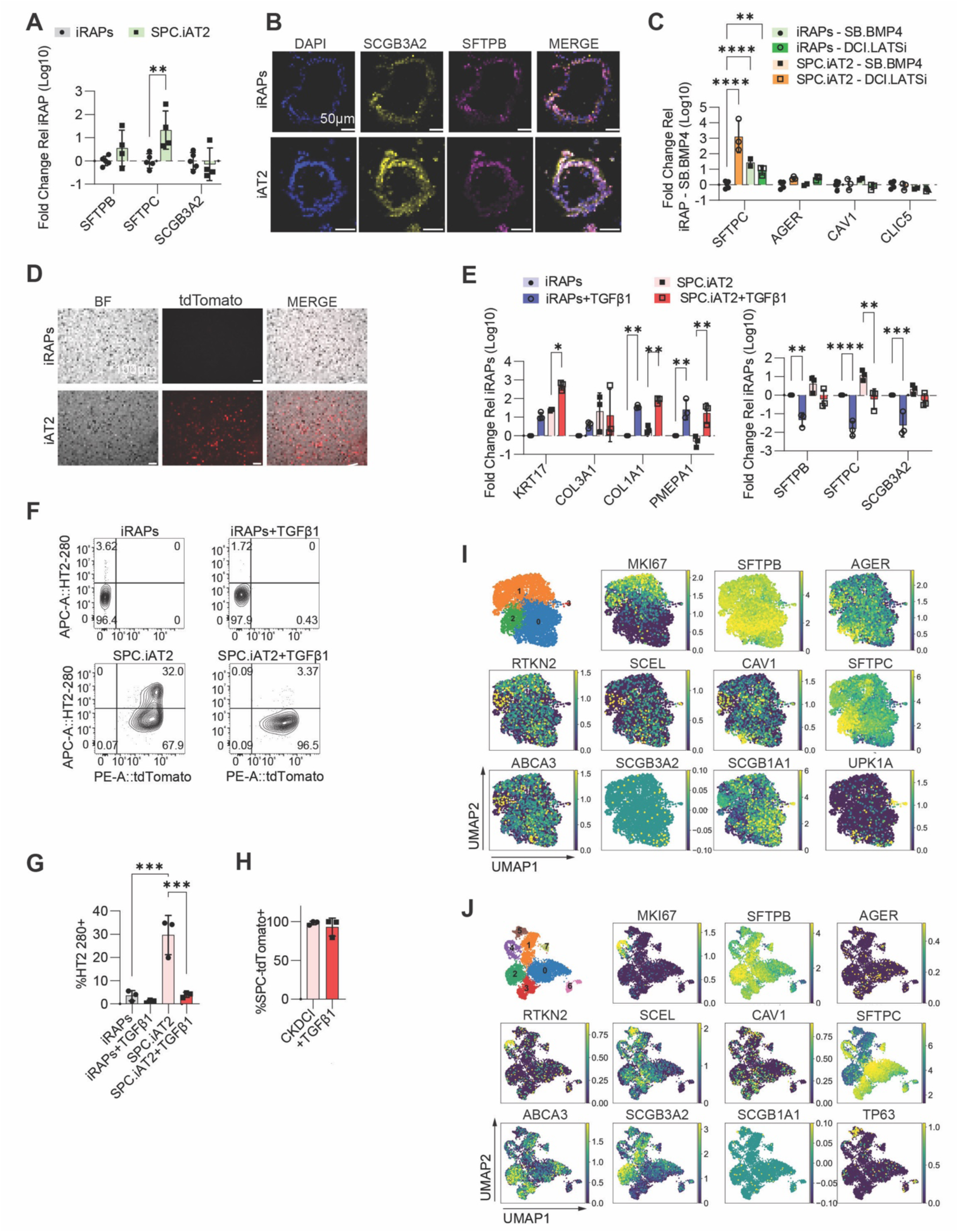
Comparison with other distal lung models. **A.** mRNA expression of TRBC markers and *SFTPC* in iRAPs and SPC-iAT2 cells. **p < 0.01, two-way ANOVA with Šidák multiple comparison test, n=5 (iRAPs) and n=4 (SPC-iAT2). **B.** Representative IF of 3 biological replicates of iRAPs (top) and SPC-iAT2 cells (bottom) stained for labelled markers. Scale bars = 50 µm. **C.** mRNA expression of iAT1 markers and *SFTPC* in iRAPs and SPC-iAT2 cells differentiated in SB-BMP4 or DCI-LATSi (right). **p < 0.01, ****p < 0.0001, two-way ANOVA with Tukey’s multiple comparison test, n=3. **D.** Representative brightfield and fluorescence microscopy images of iAT1 cells generated from iRAPs and SPC-iAT2 cells differentiated in SB-BMP4. **E.** mRNA expression of aBC markers (left) and TRBC and AT2 markers (right) in iRAPs and SPC-iAT2 cells treated with vehicle or with TGFβ1. *p<0.05, **p<0.01, ****p<0.0001, two-way ANOVA with Tukey’s multiple comparison test, n=3. **F-H.** Representative flow cytometry plot **(F)** of percent HT2-280^+^ cells **(G)** and tdTomato^+^ cells **(H)** in iRAPs and SPC-iAT2 cells treated with vehicle or with TGFβ1. ***p<0.001, one-way ANOVA with Tukey’s multiple comparison test, n=3 for HT2-280; unpaired two-sided Student’s t-test, n=3 for tdTomato. **I.** UMAP feature plots of organoids generated from ferret lung.^87^ Cluster 0 primarily expressed *SCGB1A1*, which was diffusely expressed, and with SFTPB marks TRBCs in ferrets, consistent with our observations in the ferret bleomycin model (Fig. 1). Cluster 1 expressed *MKI67*, and is therefore proliferative. The much smaller cluster 2 expressed high levels of *SFTPC*. However, some cells in this cluster co-expressed the AT1 marker, *AGER*, suggesting that they are actually ‘transitional AT2’^54^ or ‘AT0’^55^ cells, which are considered a TRBC subset.^55^ Finally, cluster 3 expressed *SCGB1A1* and *UPK1B*, and are therefore likely variant SCs, which in mice (where they are marked by *UPK3A*) possess distal regenerative capacity.^33^ **J.** UMAP feature plots of adult human distal lung organoids.^132^ AT2 cells are present (cluster 3), but other clusters correspond to *SCGB3A2^+^* TRBCs (clusters 1,2) or *SFTPC^+^SCGB3A2^+^* AT0 cells (clusters 0). A proliferating cluster 4 expressed both *SFTPB* and *RTKN2*. Cluster 6 may be consistent with cells transitioning from BCs to SC by virtue of expression of *TP63* and *SCEL* (see also Fig. 6D). The identity of clusters 6 and 7 is unclear.

**Figure S5.**
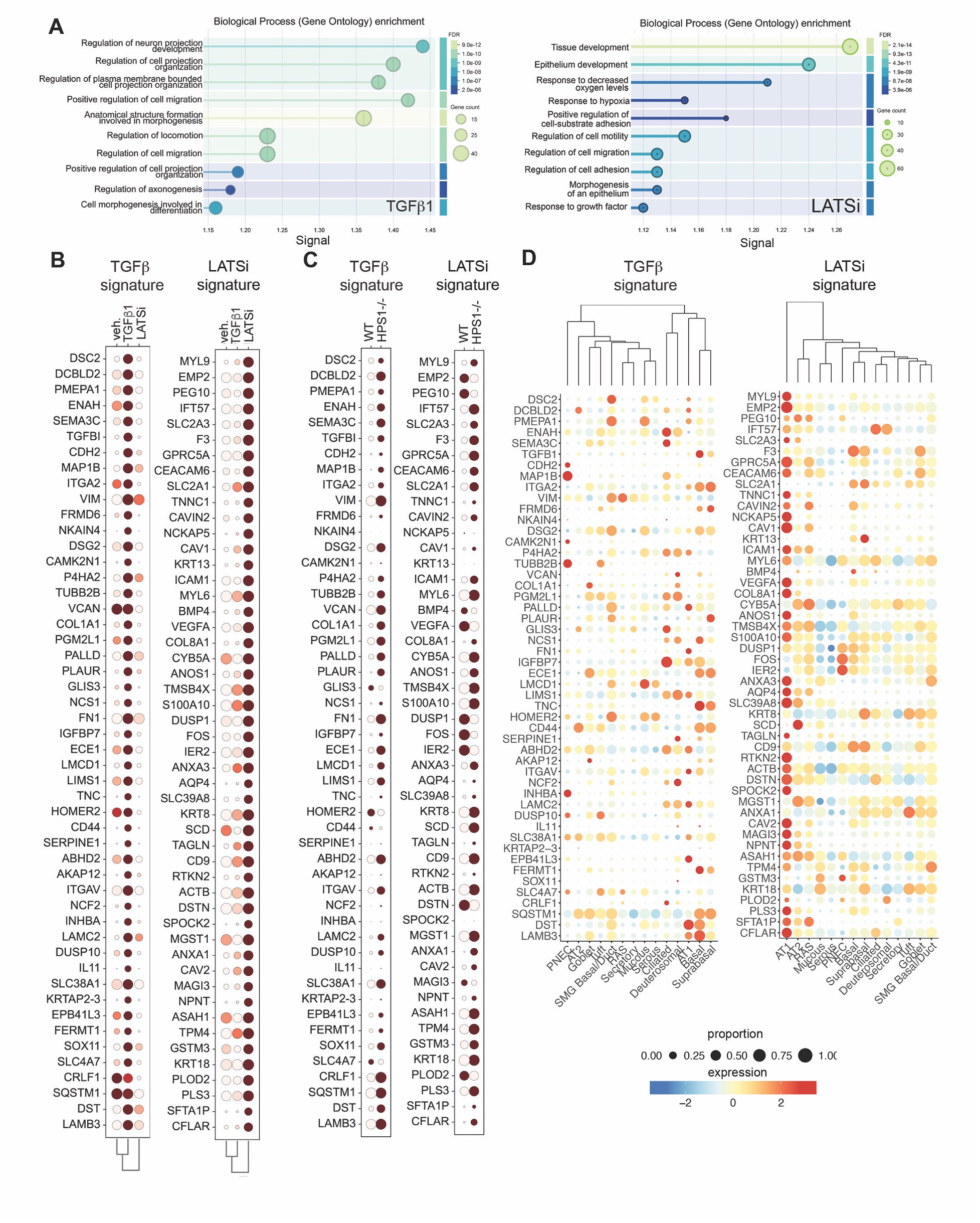
TGFβ and LATSi signatures. **A.** Gene Ontology enrichment analysis of top-200 DEGs in TGFβ1 and LATSi treated cells, respectively**. B.** Expression of TGFβ and LATSi signature genes in vehicle, TGFβ1 and LATSi-treated iRAPs. **C.** Expression of TGFβ and LATSi signature genes in WT and *HPS1^-/-^* iRAPs. **D.** Expression of genes in TGFβ and LATSi scores across all annotated epithelial cell types in the normal human lung dataset in LungMap with hierarchical clustering.

**Figure S6.**
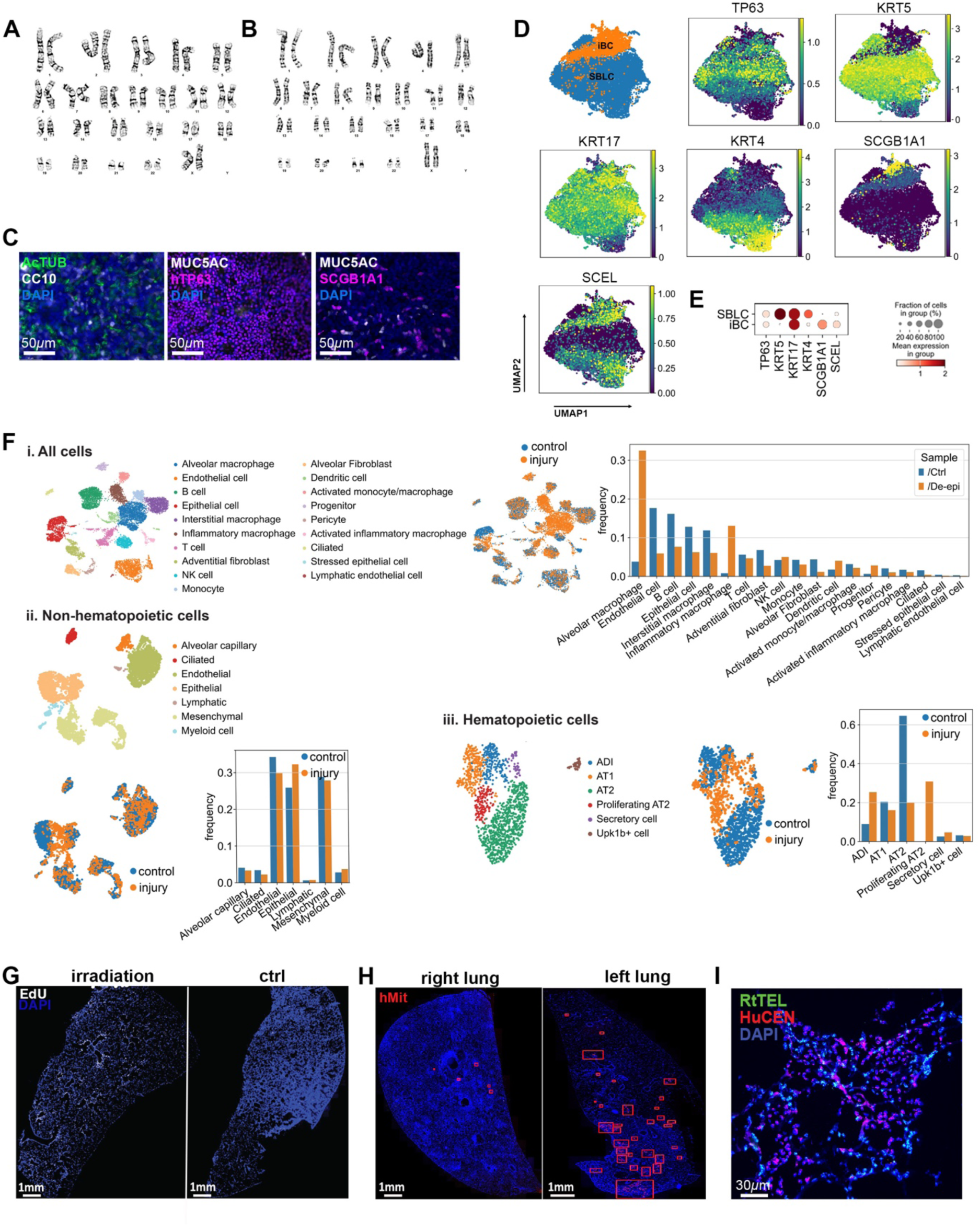
SBLC characterization and engraftment. A,B. Representative example of karyotype analysis in iPSC **(A)** and ESC-derived **(B)** SBLCs. **C.** IF of ALI cultures of SBLCs. **D.** Integrated single-cell transcriptomic analysis of iBCs ^92^ and SBLCs. **E.** Dotplot for comparative expression of select markers in iBCs and SBLCs. **F.** scRNAseq analysis of CHAPS injured rat lungs after 48h with cell types (left), samples (middle) and relative proportion of cell types (right) in the total lung population **(i)**, in the non-hematopoietic subset **(ii)** and in the epithelial subset **(iii)** in the left (injured) and right (control) lung. **G.** EdU incorporation after injury with and without irradiation. **H.** Tile scans indicating presence of hMIT+ human cells in right lung (control) and left lung (engrafted) at day 10 of the transplantation protocol. **I.** FISH for rat telomeres and human centromeres in area of rat lung engrafted with SBLCs.

**Figure S7.**
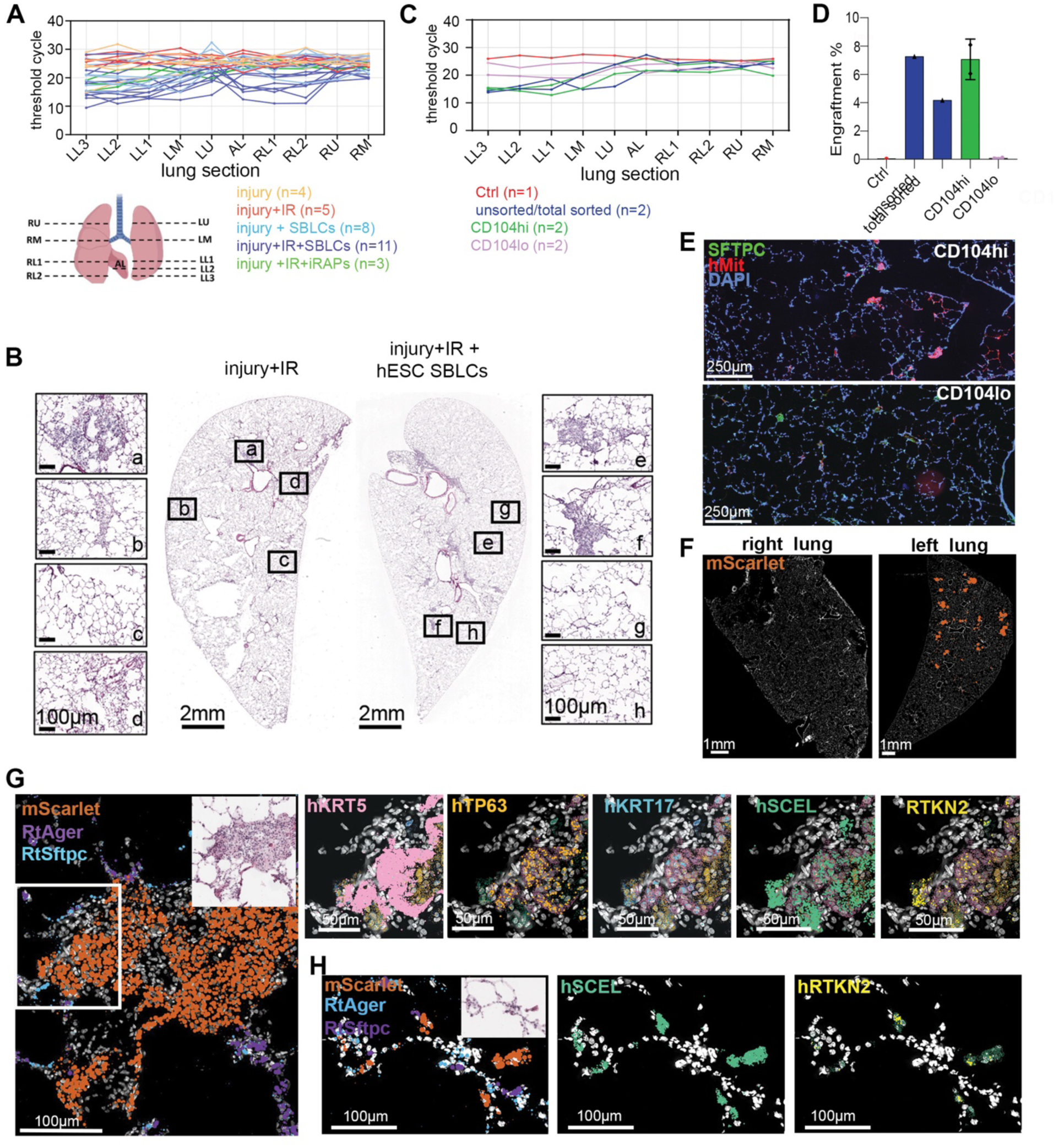
Transcriptomic analysis of engraftment. **A.** Raw DNA-qPCR data across indicated lung sections for individual experiments. **B.** H&E staining of injured and irradiated (injury + IR) and engrafted (injury + IR + hESC SBLCs) lower left lobes with representative higher magnification images. **(a, b):** inflammatory infiltrate, **c:** septal thickening, **d:** hyaline membranes, inflammatory infiltrates, **e,f:** pods, **g,h:** normal lung. **C.** Raw DNA-qPCR data across indicated lung sections for individual experiments where either CD104^hi^ of CD104^lo^ SBLC subsets were transplanted. **D.** Engraftment efficiency of CD104^hi^ of CD104^lo^ SBLC subsets as determined by ratiometric rat/human DNA qPCR. **E.** Representative example of engraftment of CD104^hi^ and CD104^lo^ SBLC subsets. **F.** Tile scan image indicating the presence of mSCARLET transcripts in right (control) and left lungs (engrafted) at day of the transplantation protocol. **G.** Representative example of Xenium spatial transcriptomics of an area containing a KRT5-pod differentiating into AT1-like cells. **H.** Representative example of Xenium spatial transcriptomics of an area containing human AT1-like cells integrated into rat alveolar structures. **I.** UMAP feature plots showing expression of indicated markers in single-cell transcriptomics of mSCARLET+ human cells isolated from engrafted rat lungs at day 10.

**Supplementary Table 1:** Datasets.

|  |  |  |
|---|---|---|
| scRNAseq nouse influenza (Niethamer et al., 2025) | GEO database | GSE262927 |
| scRNAseq mouse bleomycin (Parimon et al., 2019) | GEO database | GSM3819978 |
| scRNAseq mouse bleomycin (Koenitzer et al, 2020) | GEO database | GSE272679 |
| scRNAseq IPF (Sikkema at al., 2023) | Human Lung Cell Atlas | https://data.humancellatlas.org/hca-bio-networks/lung/atlases/lung-v1-0 |
| scRNAseq Covid (Melms et al., 2021) | GEO database | GSE171524 |
| scRNAseq pig lung homeostasis (Raredon et al., 2019). | GEO database | GSE133747 |
| WT ferret AT2 organoids (Yang et al. 2026) | GEO database | GSM9651183 |
| Distal lung organoids (Dost et al.) | GEO database | GSM8741358 |
| scRNAseq induced respiratory airway progenitors (Pezet et al., 2026) | GEO database | GSE245723 |
| scRNAseq Induced basal cells (Suzuki et al., 2021) | GEO database | GSM4223672 |
| scRNAseq SBLCs (this publication, Fig. 6D, S6D) | GEO database | GSE240574 |
| scRNAseq engrafted human cells (this publication, Fig. 7B-E, S7I) | GEO database | GSE347957 |
| Xenium engrafted rat lungs (this publication, Fig. 7A, S7G,H) | GEO database | GSE348504 |
| scRNAseq iRAPs exposed to TGFβ1 and LATSi (this publication, Fig. 5A,B, Fig. S5A,B) | GEO database | GSE240574 |

